# Offspring production of ovarian organoids derived from spermatogonial stem cells by chromatin reorganization

**DOI:** 10.1101/764472

**Authors:** Huacheng Luo, Xiaoyong Li, Geng G. Tian, Dali Li, Changliang Hou, Xinbao Ding, Lin Hou, Qifeng Lyu, Yunze Yang, Austin J. Cooney, Wenhai Xie, Ji Xiong, Hu Wang, Xiaodong Zhao, Ji Wu

## Abstract

Fate determination of germline stem cells remains poorly understood at the chromatin structure level^1,2^. Here, we demonstrate successful production of offspring from oocytes transdifferentiated from mouse spermatogonial stem cells (SSCs) with tracking of transplanted SSCs in vivo, single cell whole exome sequencing, and in 3D cell culture reconstitution of the process of oogenesis derived from SSCs. Furthermore, we demonstrate direct induction of germline stem cells (iGSCs) differentiated into functional oocytes by transduction of *H19, Stella*, and *Zfp57* and inactivation of *Plzf* in SSCs after screening with ovarian organoids. Using high throughput chromosome conformation, we uncovered extensive chromatin reorganization during SSC conversion into iGSCs, which was highly similar to female germline stem cells. We observed that although topologically associating domains were stable during SSC conversion, chromatin interactions changed in a striking manner, altering 35% of inactive and active chromosomal compartments throughout the genome. These findings have important implications in various areas including mammalian gametogenesis, genetic and epigenetic reprogramming, biotechnology, and medicine.

## Main

Cell fate decisions, which require key gene regulation, remain poorly understood at the chromatin structure level. Although three-dimensional chromatin architectures of mouse gametes were recently reported, how they affect fate decisions of germline stem cells remains to be explored^1-3^.

To characterize spermatogonial stem cells (SSCs), we firstly isolated by magnetic activated cell sorting (MACS) using an anti-Thy-1 antibody after two step enzymatic digestion of the testes from 6-day-old pou5f1 (also known as Oct4, a germ cell-specific transcriptional factor)/GFP transgenic×C57BL/6 F1 hybrid mice^4^. Then, the isolated SSCs were purified for GFP-positive SSCs by fluorescence-activated cell sorting (FACS) (Extended Data Fig. 1a). The purified SSCs were maintained on SIM mouse embryo derived thioguanine- and ouabain-resistant (STO) feeder layers (see METHODS) (Extended Data Fig. 1b). After 3–5 days of culture, SSCs expanded into clusters (Extended Data Fig. 1c). Next, we determined the expression patterns of *Oct4, Mvh* (mouse vasa homologue, expressed exclusively in germ cells), *c-Ret*^5^, *Plzf*^6^, *Rex-1*^7^, *Utf1*^8^, *Esg-1* (also known as *DPPA5*)^9^, *Stra8*^10^, *Sox2*^11^, and *Nanog*^12^. Reverse transcription-polymerase chain reaction (RT-PCR) and immunocytochemical analyses showed that SSCs expressed *Oct4, Mvh, c-Ret, Plzf, Rex-1, Utf1, Esg-1*, and *Stra8*. Cytogenetic analysis by treatment with colchicines followed by G-band staining demonstrated a normal karyotype (40, XY) in the metaphase spreads of examined SSCs (Extended Data Fig. 1d–j). To verify the characterization of SSCs, we compared the global expression profiles of SSCs and embryonic stem cells (ESCs) using microarrays. Gene expression profiles by scatter plots showed a significant difference between SSCs and ESCs (Extended Data Fig. 1k, n=3). More than two thousand genes (2251) were differentially expressed between SSCs and ESCs, including pluripotency-related genes *Dppa4, Fgf4, Nanog, Sox2*, and *Klf4* (Extended Data Fig. 1k, l, fold change>2, P<0.05, t-test) and SSC-related genes *Zbtb16(or Plzf), Gfra1, Tex18, Piwil2*, and *Dazl* (Extended Data Fig. 1k, l, fold change>2, P<0.05, t-test). Therefore, these results demonstrated that SSCs had their apparent original property rather than a pluripotent identity. To determine the imprinting pattern of SSCs, differentially methylated regions (DMRs) of two paternal (*H19* and *Rasgrf1*) and two maternal (*Igf2r* and *Peg 10*) imprinted regions were examined in SSCs and ESCs by bisulfite genomic sequencing. In SSCs, paternally imprinted regions (Extended Data Fig. 1m, o) were methylated, while maternally imprinted regions were not methylated (Extended Data Fig. 1n, p); this indicated an androgenetic imprinting pattern that was different from that of ESCs.

For investigating SSC fate determination in the mouse ovary, pou5f1/GFP transgenic mouse SSCs cultured for 3–5 days were directly transplanted into the ovaries of premature ovarian failure (POF) mice (see METHODS). Phosphate buffered saline (PBS) was injected into the ovaries of POF recipients as a control. For the positive control, female germline stem cells (FGSCs) from pou5f1/GFP transgenic mice were also transplanted into the ovaries of POF mice (see METHODS). At 8 weeks post-transplantation, recipient ovaries including positive control ovaries were collected and evaluated for morphology and GFP expression. Histological analysis showed that recipient ovaries injected with cells contained numerous oocytes at all stages of development, including GFP-positive oocytes (Fig. 1a I, III, IV and Extended Data Fig. 2a-e). Furthermore, DNA fluorescence in situ hybridization (FISH) analysis showed the presence of the *Sry* gene in oocytes from recipient ovaries (Fig. 1b). For confirmation, single cell whole exon sequencing was used. The results demonstrated that the germinal vesicle (GV) oocytes from recipient ovaries were derived from transplanted SSCs (Fig. 1c). Mature oocytes from recipient ovaries were then collected for karyotype analysis. The results showed that some mature oocytes had the karyotype of 20, Y (Fig. 1d I–V). PCR analysis of DNA fragment *Sry* confirmed that some mature oocytes contained a candidate of the Y chromosome (Fig. 1d VI). However, control ovaries consisted of stromal and interstitial cells as well as atretic follicles (Fig. 1a II). These results indicate that XY oocytes were regenerated in POF females by transplantation of SSCs.

**Fig. 1:**
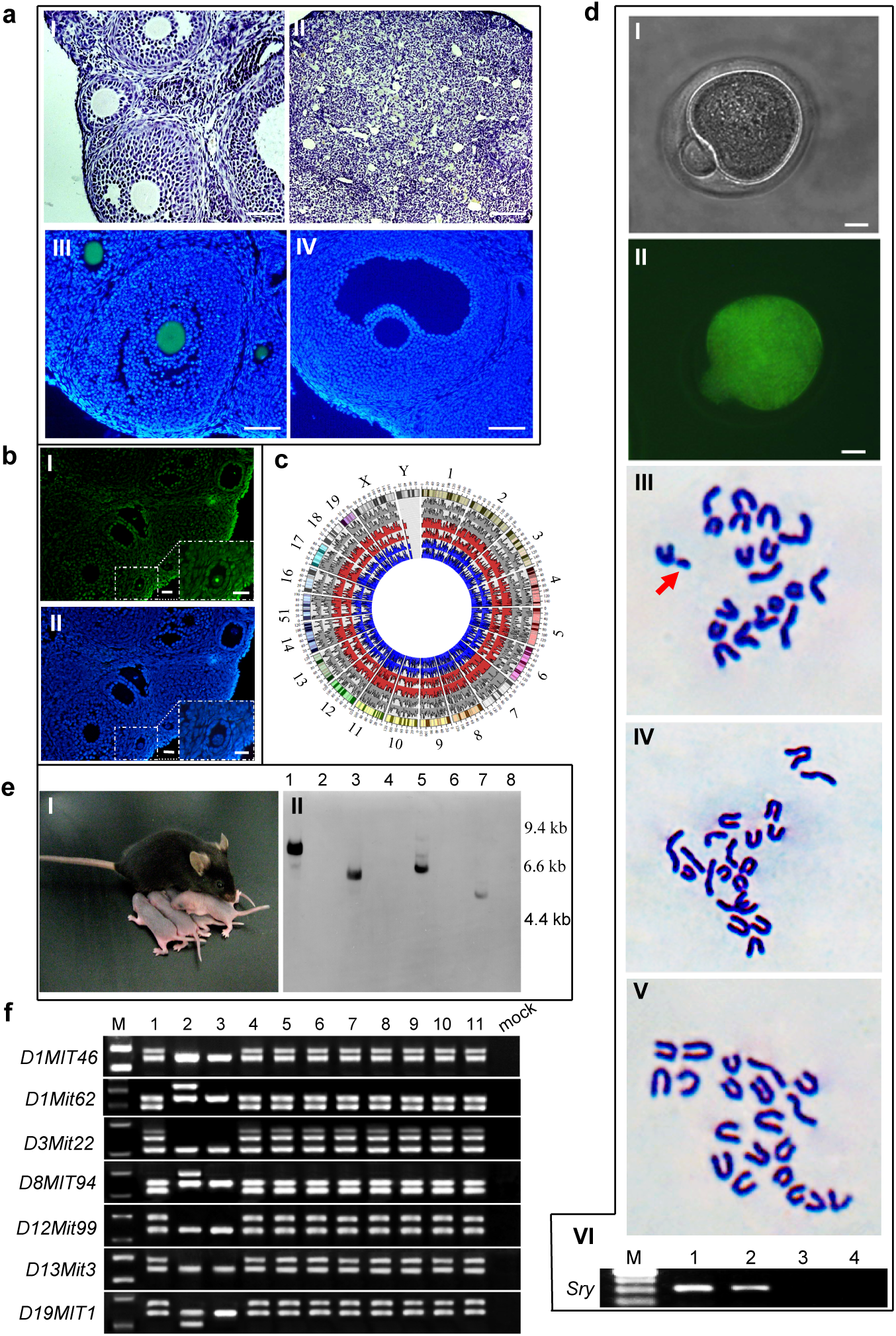
SSCs transdifferentiate into oocytes in the ovaries of POF recipients and GFP-expressing offspring are generated from the transplanted SSCs from pou5f1/GFP transgenic mice. **a**, SSCs were transplanted into the ovaries of POF recipient mice. I, II, Representative morphologies of the ovaries from recipients with (I) or without (II) SSC transplantations. III, Follicles containing GFP-positive (green) oocytes in recipient ovaries at 8 weeks after transplantation of pou5f1/GFP transgenic SSCs. IV, Oocytes in a wild-type ovary without a GFP signal. **b**, DNA fluorescence *in situ* hybridization for SRY. SRY was only localized in oocytes (green) derived from SSCs in ovary (I). Nuclei were counterstained with DAPI (blue) (II). **c**, Circos plot showing the coverage from the single cell exon sequencing as a histogram. Grey represents FGSCs; Red represents GV oocytes derived from SSCs in the ovary; Blue represents SSCs. **d**, Karyotype analysis of mature oocytes from POF recipient ovaries at 2 months after pou5f1/GFP transgenic SSC transplantation. I, II, Representative morphologies of mature oocytes derived from pou5f1/GFP transgenic SSCs (I) emitting GFP fluorescence (II) under UV light. III–V, Cytogenetic analysis by G-band staining showing that some mature oocytes from SSCs had a karyotype of 20, Y. III: An example of 20, Y in mature oocytes derived from pou5f1/GFP transgenic SSCs. Arrow indicates the Y chromosome. IV: Example of 20, X in mature oocytes derived from pou5f1/GFP transgenic SSCs. V: Representative karyotype (20, X) of wild-type mature oocytes. VI, PCR analysis of *Sry*. M, 100 bp DNA marker; lane 1, SSCs; lane 2, mature oocytes derived from pou5f1/GFP transgenic SSCs; lane 3, wild-type mature oocytes; lane 4, mock. **e**, Example of offspring from POF recipient mice transplanted with pou5f1/GFP transgenic SSCs (I) and an example of a Southern blot of tail DNA (II). Genomic DNA was digested with *Eco*RI. Marker sizes are indicated to the right of the blot. Lanes 1, 3, 5, and 7: transgenic mice; lanes 2, 4, 6, and 8: wild-type mice. **f**, SSLP analysis of parents and their offspring mice through SSLP markers. M: DNA marker; lane 1: donor SSCs; lane 2: female recipients (POF); lane 3: mated males (C57BL/6); lanes 4–11: offspring from eight corresponding recipients females. Scale bars, 50 μm (**a** I, III, IV), 100 μm (**a** II), 25 μm (**b** I, II), 10 μm (**d** I, II).

To examine whether XY oocytes derived from SSCs could produce offspring, POF recipients were mated with wild-type C57BL/6 adult males at 35 days after cell transplantation or PBS injection (control)^13,14^. Control recipients were not fertile (*n* = 9). All POF recipients produced offspring (n=8, Fig. 1e I) with more males than females per litter (male:female, 1.95:1.00). One hundred and sixteen of the 130 offspring were alive with a normal phenotype as well as fertile. Fourteen of the 130 offspring died at 1–6 weeks after birth. The offspring were examined for the presence of GFP transgenes by Southern blot analyses (Fig. 1e II). Sixty-four of the 130 F1 progeny were heterozygous for the GFP transgene. Furthermore, simple sequence length polymorphism (SSLP) analysis was performed with SSLP markers to confirm that the offspring were derived from transplanted SSCs. The offspring from eight recipients (see above) were distinct from POF (see METHODS) or C57BL/6 mice-their parents (POF mice and mated male); however, they had exactly the same profiles as the SSCs from which they were derived (Fig. 1f). After analysis of the methylation status, five of the 20 offspring demonstrated abnormal methylation patterns. Two offspring that did not survive showed high methylation in *Peg 10* with an increase of 21.46%±2.66% and 28.16%±2.87% compared with the control. The remaining three demonstrated that *H19* was highly methylated and increased by 16.66%±1.48% and 14.23%±1.38% or had low methylation with a decrease of 14.94%±1.26% compared with the control. Moreover, five out of six positive controls (see METHODS) with FGSC transplantation were fertile with approximately equal numbers of males and females per litter (male:female, 1.02:1.00), and their offspring showed no abnormal phenotype. Fifty of the 101 offspring were heterozygous for the GFP transgene (Extended Data Fig. 2f, g). Although approximately 11% of the offspring were abnormal, these results suggest that the XY oocytes derived from SSCs can produce offspring in previously sterile recipients and generate transgenic progeny. Control adult mice that received PBS injections into their ovaries did not produce transgenic offspring.

For understand how the SSCs transdifferentiated into oocytes in recipient ovaries, pou5f1/GFP transgenic mouse SSCs cultured for 3 days were directly transplanted into the ovaries of POF mice, and then monitored by confocal laser scanning microscopy. At 2 hours after SSC transplantation, the SSCs were observed in ovaries of recipient mice, indicating that the SSCs had been successfully transplanted into the mouse ovary (Fig. 2a). The transplanted cells were found to migrate toward the edge of the ovarian cortex at 2 days post-transplantation (Fig. 2a). At 3 days after transplantation, the cells continuously migrated toward the edge of the ovarian cortex and some of them reached the edge (Fig. 2a). When the transplanted cells had been in the ovary for 4 days, all of them had migrated into the edge of the ovarian cortex (Fig. 2a). Five days post-transplantation, transplanted cells settled in the edge of the ovarian cortex and began to transdifferentiate into early primary oocytes (Fig. 2a, b). At 6–15 days after transplantation, the transplanted cells continued to transdifferentiate into oocytes at various stages of development (Fig. 2a). This was confirmed by dual immunofluorescence analysis of the expression of MVH and GFP in transplanted cells (Fig. 2c).

**Fig. 2:**
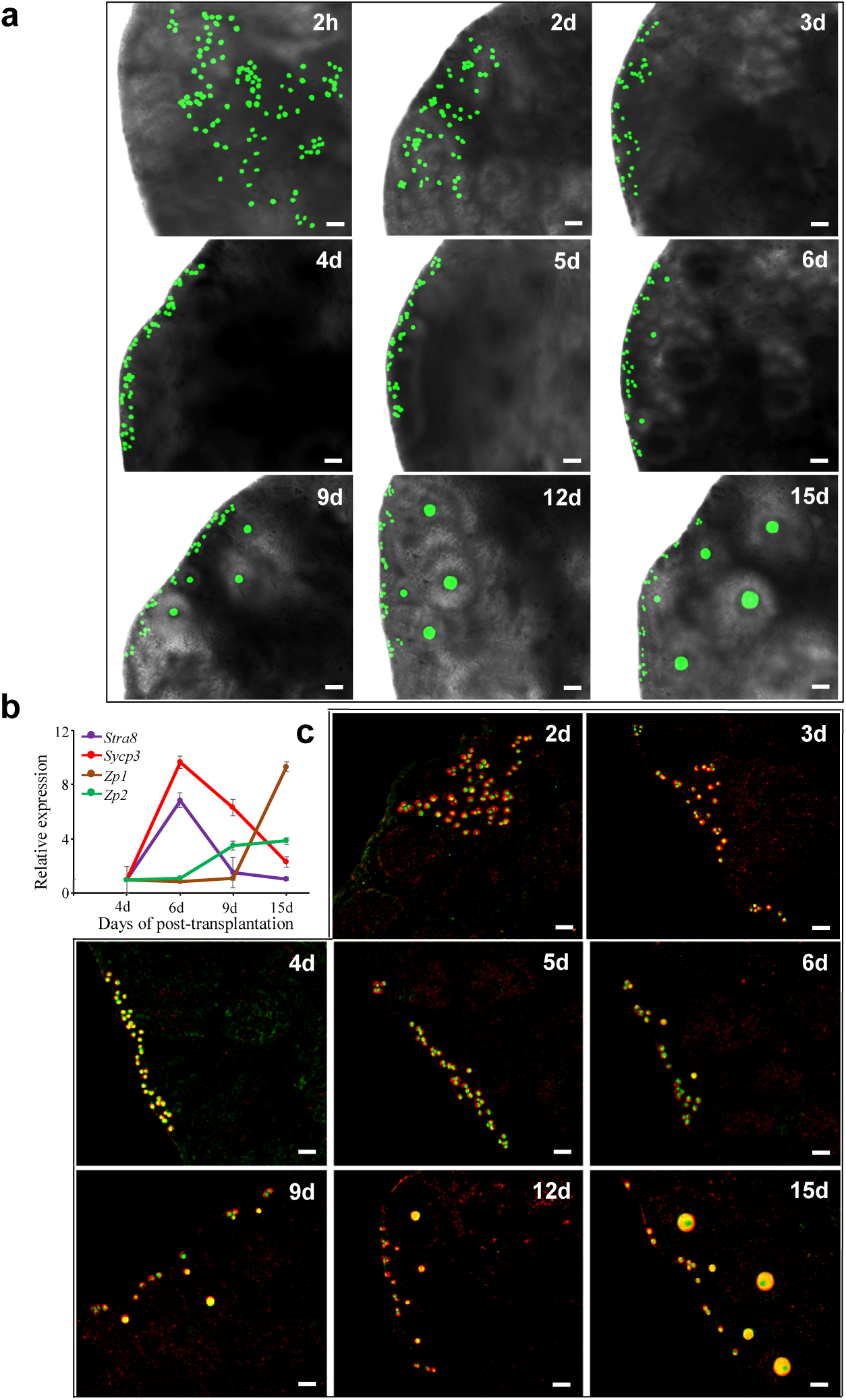
Tracking of transplanted SSCs in recipient ovaries. **a**, Transplanted SSCs from pou5f1/GFP transgenic mice were monitored by confocal laser scanning microscopy at 2 hours, and 2, 3, 4, 5, 6, 9, 12, and 15 days after transplantation into recipient ovaries. **b**, Gene expression dynamics during oogenesis in transplanted cells at 4, 6, 9, and 15 days after transplantation. **c**, Dual immunofluorescence analysis of MVH and GFP expression in transplanted cells at 2, 3, 4, 5, 6, 9, 12, and 15 days after transplantation. Scale bars, 50 μm.

To explore the mechanism of SSC fate determination when the SSCs were transplanted into the recipient ovary, we performed bisulfite sequencing to analyze the methylation status of these transplanted cells, mainly the differentially methylated regions (DMRs) of paternally imprinted gene *H19* and maternally imprinted gene *Peg10*. It is noteworthy that no obvious change of methylation levels, including the maternally or paternally imprinting gene, was observed at 2 hours after SSC transplantation (Extended Data Fig. 3a–d). At 3–4 days after transplantation, methylation levels of *H19* were reduced gradually to 68.3% and 60.8%, with a further reduction to 37.9% and 23.8% at 5–6 days, suggesting that the bulk of methylation erasure occurred at 5–6 days. The low levels of methylation were present at 9 days and persisted to 15 days (Extended Data Fig. 3a, c). In contrast, the maternally imprinted gene *Peg10* showed evidence of robust de novo methylation with an increase to 45.7% methylation at 3 days and further increase to 78% methylation at 6 day, indicating that the bulk of methylation establishment occurred at 6 days. The high methylation levels were maintained from 9 to 15 days (Extended Data Fig. 3b, d). These results suggested that the maternal DNA methylation pattern was directly constructed during SSC development in recipient ovaries, and that the SSCs did not dedifferentiate into PGCs.

Furthermore, we determined expression patterns of imprinted genes (*H19, Grb10, Gtl2, Rasgrf1, Peg10, Igf2r*, and *Snrpn*) and important transcription factor genes (*Plzf, Stella or Dppa3, Zfp42, Zfp57*, and *Nanos2*) during transdifferentiation of the transplanted cells into oocytes in recipient ovaries based on data from our previous studies^15,16^. After comparing the expression of imprinted genes in the cultured SSCs (0 hour) and transplanted SSCs at 2 hours to 15 days post-transplantation, we observed that paternally imprinted genes (*H19, Grb10, Rasgrf1*, and *Gtl2*) and transcription factor genes (*Stella* and *ZFP57*) were gradually upregulated, especially at 3 and 6 days with a further increase from 9 to 15 days (Extended Data Fig. 3e). In contrast, along with transdifferentiation into oocytes of the SSCs in recipient ovaries, the expression levels of these maternally imprinted genes (*Peg 10, Igf2r*, and *Snrpn*) and transcription factor genes (*Plzf, Zfp42*, and *Nanos2*) underwent obvious reductions at 5–6 days with a continuous decrease from 9 to 15 days after transplantation (Extended Data Fig. 3e), suggesting establishment of the maternal imprinting pattern.

Based on the above results, we further screened for the critical imprinted genes and transcription factor genes required for SSC conversion. The spatial organization of the mammal genome is known to play an important role in the regulation of gene expression^17^. Therefore, we used in situ high throughput chromosome conformation capture (Hi-C) to further screen for the critical genes and found 6.28 billion unique read pairs in SSCs and FGSCs. The compartment status was divided into two groups, compartment A and B. By comparing A/B compartment statuses and chromatin loops between SSCs and FGSCs (Extended Data Fig. 4, Supplementary Table 1-3), we established combinations of six genes (6Gs), including imprinted genes, *H19* and *Rasgrf1*, and transcription factor genes *Stella, Zfp57, Zfp42*, and *Plzf*. After overexpressing *Stella, H19, Zfp57*, and *Rasgrf1* and knockdown of *Plzf* and *Zfp42* in SSCs, the cells converted to induced germline stem cells (or induction of germline stem cells, iGSCs) with a maternal imprinted pattern (Extended Data Fig. 5, Extended Data Fig. 6a) and formed ovarian organoids when three-dimensional (3D) co-cultured with somatic cells from the fetal ovary for 2 weeks, modifying the method previously described in ref^18^ (Extended Data Fig. 6b I-II). Upon withdrawal of *Rasgrf1* from the 6Gs, we found that the 3D co-cultured cells still formed ovarian organoids (Extended Data Fig. 6b III). For the remaining five genes (5Gs), removal of *Zfp42* further promoted the formation of the organoids (Extended Data Fig. 6b IV). However, removal of any factor from the four genes (4Gs, *Stella, H19, Zfp57*, and *Plzf*) led to the failure to form the organoids (Extended Data Fig. 6b V–VIII).

The morphology of iGSCs induced by the overexpression of *Stella, H19*, and *Zfp57* and the inactivation of *Plzf* was similar to that of FGSCs (Extended Data Fig. 7a). Furthermore, the iGSCs expressed *Stella, Mvh, Fragilis, Dazl*, and *Oct4* with a maternal imprinted pattern (Extended Data Fig. 7a, b, Extended Data Fig. 6c). For confirmation, we performed genome-wide DNA methylation analysis in SSCs, iGSCs, and FGSCs by MeDIP-seq. A total of 38.7 million reads, yielding 467,163 DNA methylationsites (peaks) in three kinds of cell populations were generated. We observed widespread variation in terms of DNA methylation during SSC transition into iGSCs (Extended Data Fig. 7c). Subsequently, we performed pair-wide correlation analysis of the MeDIP data sets from SSCs, iGSCs, and FGSCs. We found that the overall DNA methylation pattern of iGSCs was similar to that of FGSCs (r = 0.79), but it was less similar to that of SSCs (r = 0.59). Such a trend was evidenced more clearly by individual regions of interest. For example, at the maternally imprinted region *Igf2r*, the DNA methylation signal was relatively low in the *Igf2r* promoter of SSCs. It increased remarkably and appeared to be almost at the same level in iGSCs and FGSCs (Extended Data Fig. 7c, d). A similar phenomenon was also observed at the promoter region of *Nr0b1*, the gene encoding the orphan nuclear receptor and required for development of male characteristics in mice^19^. These observations suggest that DNA methylation contributed to the SSC transition. Next, we compared global gene expression profiles among SSCs, iGSCs and FGSCs by RNA sequencing. A total of 380,702,626 raw reads were generated. We detected expression of 18229, 19755, and 18978 out of 24550 genes in SSCs, iGSCs, and FGSCs, respectively. On average, 77% of the known mouse genes were expressed in the sampled SSCs, iGSCs, and FGSCs. Hieratical clustering was performed, and the results indicated that iGSCs were clustered with FGSCs, but separated from SSCs, suggesting that the global gene expression profile of iGSCs was similar to that of FGSCs (Extended Data Fig. 7e, f). Among these genes, *Igf2r*, a maternal imprinted gene, showed high expression in SSCs and low level expression in iGSCs and FGSCs (Extended Data Fig. 7f), which was consistent with the results from the analysis of genome-wide DNA methylation in SSCs, iGSCs, and FGSCs.

Hi-C interaction maps provide information on multiple hierarchical levels of genome organization^20^. To understand how genome organization is involved in SSCs transition to iGSCs, we also performed Hi-C experiments using two biological replicates of iGSCs, generating a total of 2.96 billion unique read pairs. The Hi-C data analysis showed the high order chromatin organization of the whole genome in SSCs, iGSCs, and FGSCs (Fig. 3a–c, Extended Data Fig. 9). To examine the characteristics of their chromatin organization, we analyzed the pattern of compartment A/B in SSCs, iGSCs, and FGSCs. We found a large degree of spatial plasticity in the arrangement of the A/B compartments or redistribution of the spatial organization of their genomes during SSC transition into iGSCs with 35% of the genome switching compartments. Furthermore, we found that the regions that changed their A/B compartment status corresponded to a single or series of topologically associated domains (TADs), suggesting that TADs are the units of dynamic alterations in chromosome compartments (Fig. 3b). Interestingly, we observed that iGSCs and FGSCs were highly similar in their status of A/B compartments compared with SSCs (Fig. 3c). For SSC transition into iGSCs, genes that changed from compartment B to A tended to show higher expression, while genes that changed from A to B tended to show reduced expression (Fig. 3d). Moreover, we identified 4353 genes with co-variation between compartment switching and gene expression. For example, at the compartment B region, *Dppa3* expression was relatively low in SSCs. It increased remarkably and appeared to be almost at the same level in iGSCs and FGSCs when changing from compartment B to A (Fig. 3e). DNA methylation that changed from compartment B to A also tended to show a reduced signal, whereas DNA methylation that changed from A to B tended to show higher signal (Fig. 3f). Take together, these results demonstrate that, when SSCs convert to iGSCs, there is a high degree of plasticity in A and B compartments, corresponding changes in gene expression, indicating that the A and B compartments have a contributory to cell type-specific patterns of gene expression.

**Fig. 3:**
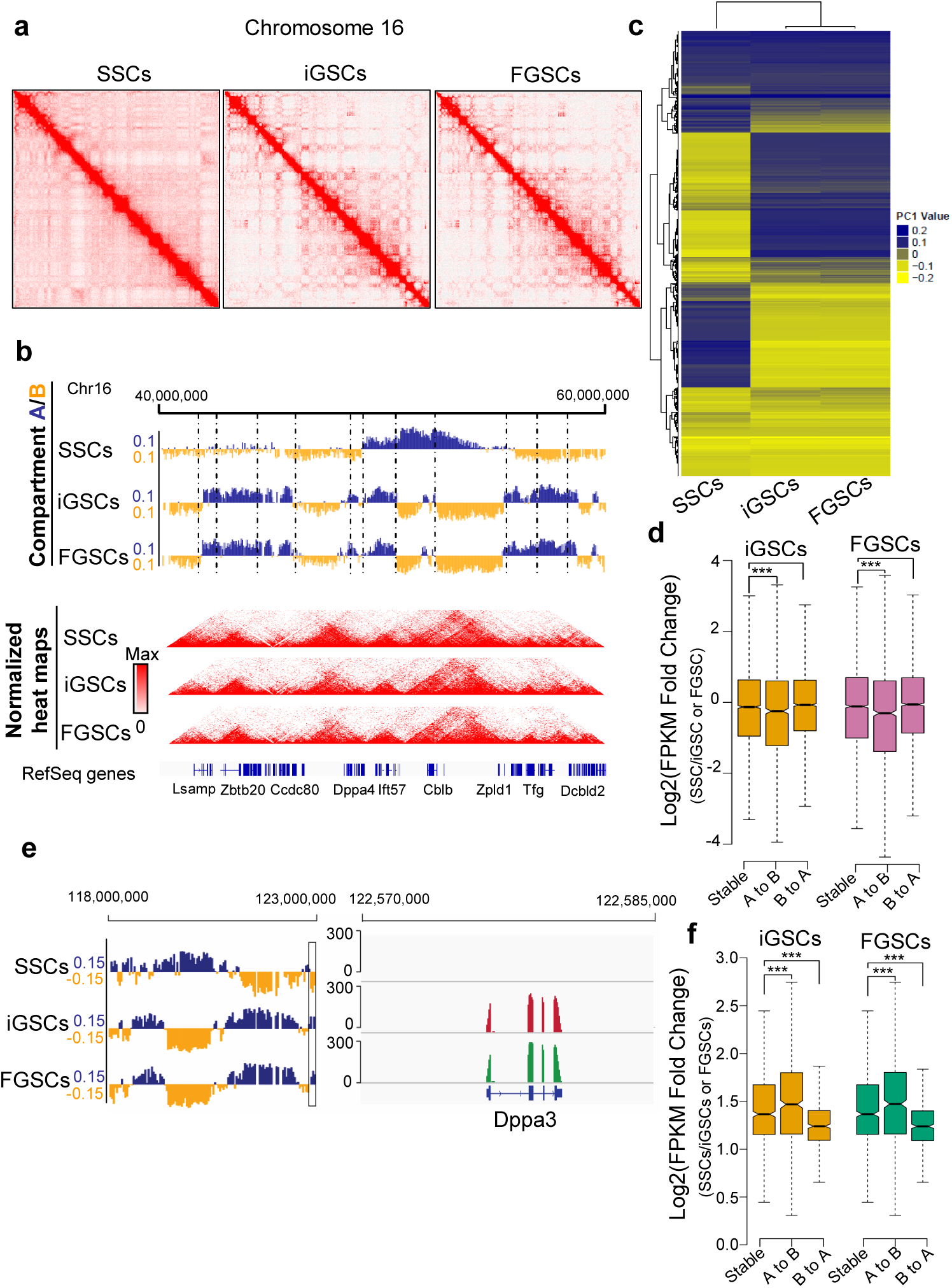
Reorganization of the chromosome structure during SSC conversion to iGSCs. **a**, Contact matrices from chromosome 16 in SSCs, iGSCs, and FGSCs. **b**, First principal component (PC1) value and normalized Hi-C interaction heat maps at a 40kb resolution in SSCs, iGSCs, and FGSCs. The PC1 value was used to indicate the A/B compartment status, where a positive PC1 value represents the A compartment (blue) and a negative value represents the B compartment (yellow). Dashed lines indicate TAD boundaries in SSCs. **c**, Hieratical clustering of PC1 values for the A/B compartment status in SSCs, iGSCs, and FGSCs. **d**, Expression of genes that changed compartment status (“A to B” or “B to A”) or remained the same (“stable”) compared with SSCs (P-value by Wilcoxon’s test). **e**, IGV snapshot of Dppa3 (Stella) showing concordance between its expression and PC1 values. **f**, Relative MeDIP-seq signal that changed compartment status (“A to B” or “B to A”) or remained the same (“stable”) compared with SSCs (P-value by Wilcoxon’s test). ***p<0.0001.

Developmental feature of ovarian organoids from FGSCs and iGSCs were explored. At 2 weeks of 3D co-culture with the germline stem cells and somatic cells from the fetal ovary, ovarian organoids were generated in FGSC and iGSC groups. The organoids were completely filled with follicles which possessed oocytes. In contrast, ovarian organoids were not formed in the SSC group. When the ovarian organoids were 3D co-cultured for 3 weeks, the follicles grew obviously (Fig. 4a). After these follicles were 3D cultured individually for 2 weeks (see METHODS), a large number of immature oocytes were obtained from the cultured follicles. Moreover, Oct4-EGFP cells were detectable in iGSC and FGSC groups at 3 days of 3D co-culture. A number of EGFP-positive oocytes were observed in ovarian organoids after 3 weeks of 3D co-culture (Fig. 4a). After 2-3 weeks of 3D co-culture, however, the EGFP expression became weak (Fig. 4a).

**Fig. 4:**
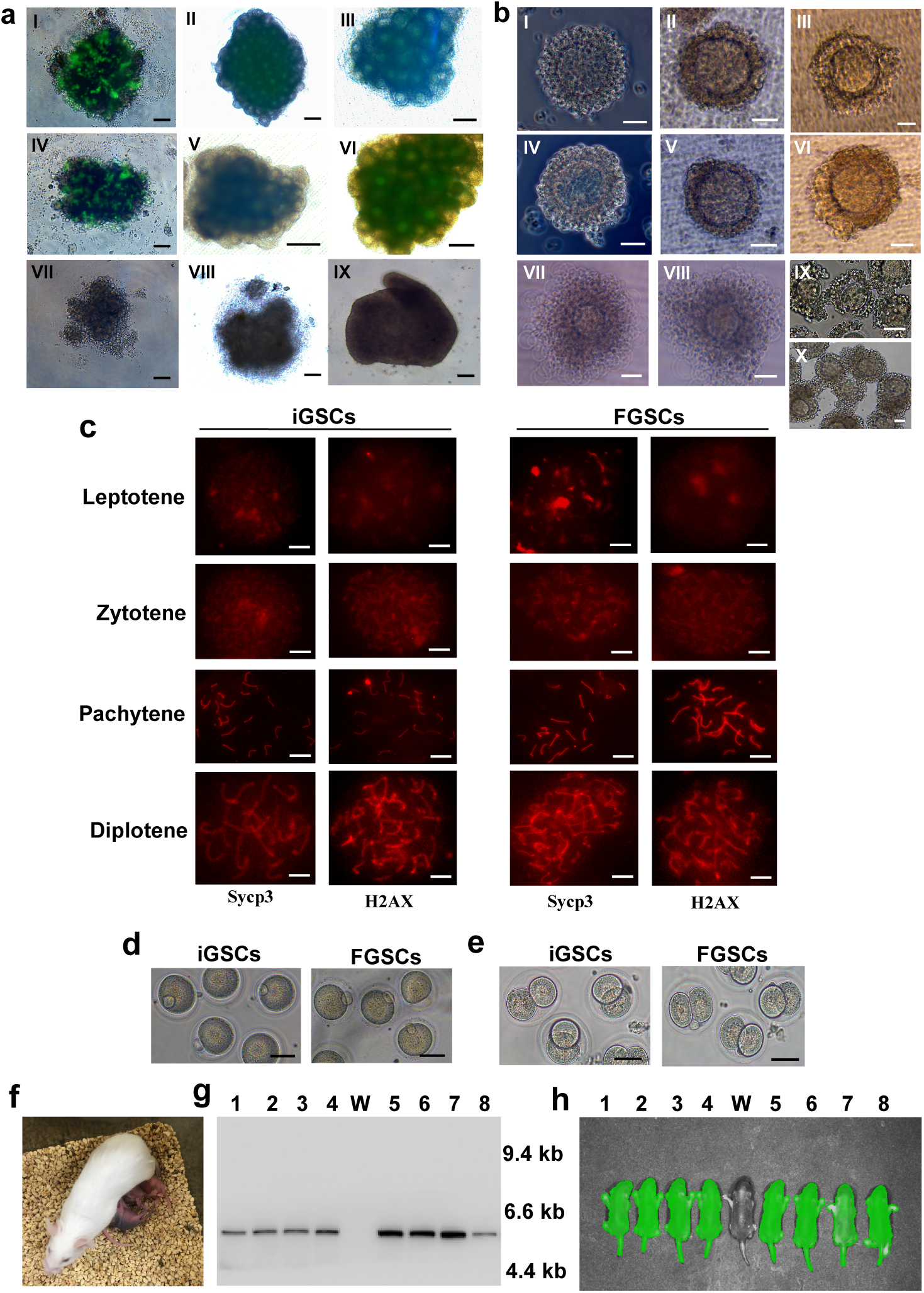
Offspring production of ovarian organoids derived from iGSCs and FGSCs. **a**, Ovarian organoid formation and development. Representative ovarian organoids with a merge of bright field and fluorescence. I–III, Ovarian organoids or co-cultures with somatic cells of gonad and iGSCs at 3 days (I), 2 weeks (II), and 3 weeks (III). IV–VI, Ovarian organoids or co-cultures with somatic cells of gonads and FGSCs at 3 days (IV), 2 weeks (V), and 3 weeks (VI). VII–IX, Images of aggregates formed by somatic cells of gonads and SSCs at 3 days (VII), 2 weeks (VIII), and 3 weeks (IX**). b**, Follicle growth in vitro. I–III, VII, Representative follicles isolated from ovarian organoids formed by somatic cells of gonads and iGSCs at 0 days (I), 2 days (II), 7 days (III), and 11 days (VII). IV–VI, VIII, Representative follicles isolated from ovarian organoids formed by somatic cells of gonads and FGSCs at 0 days (IV), 2 days (V), 7 days (VI), and 11 days (VIII). IX, X, Cumulus-oocytes complexes derived from iGSCs (IX) and FGSCs (X) before in vitro maturation. **c**, Representative views of each stage of meiotic prophase I during ovarian organoid development after stained with anti-Sycp3 and -H2AX antibodies. **d**, Mature oocytes derived from iGSCs or FGSCs after in vitro maturation. **e**, Two-cell embryos derived from iGSCs or FGSCs after in vitro fertilization. **f**, Representative offspring derived from iGSCs or FGSCs. **g**, Offspring were identified by Southern blotting. Lanes 1–4, offspring derived from iGSCs, lane W, wild-type mice, lanes 5–8, offspring derived from FGSCs. **f**, Offspring were identified by fluorescence. Lanes 1–4, offspring derived from iGSCs, lane W, wild-type mice, lanes 5–8, offspring derived from FGSCs. Scale bars, 100 μm (**a**), 20 μm (**b** I-VI), 40 μm (**b** VII-VIII), 50 μm (**b** IX-X, **d, e**), 5 μm (**c**).

For comparison between iGSC and FGSC groups, 905 immature oocytes from the iGSC group were obtained from 27 ovarian organoids in 8 cultures with 33.52±4.27 immature oocytes per organoid. In the FGSC group, 1140 immature oocytes were obtained from 36 ovarian organoids in 12 experiments with 31.67±3.86 immature oocytes per organoid. No difference in immature oocytes per organoid was observed in both groups (Fig. 4b).

To evaluate oogenesis during the ovarian organoid development, immunofluorescence analysis of SCP3 and H2AX was performed. Furthermore, the gene expression profiles during iGSC or FGSC differentiation were analyzed by qRT-PCR. The results showed progression of meiotic prophase I from 7 to 21 days of the germ cell differentiation in both groups (Fig. 4c). The expression dynamics of genes involved in oogenesis were similar between both groups (Extended Data Fig. 10).

After in vitro maturation for 17-20 hours, 48.9% and 51.3% of these immature oocytes reached mature oocytes in iGSC and FGSC groups, respectively. In addition, no difference in the number of mature oocytes was observed between the two groups (Fig. 4d, Extended Data Fig. 11a).

To determine whether these mature oocytes could develop into offspring following in vitro fertilization, embryo culture and transfer into pseudopregnant ICR females were performed. The fertilization rate of the iGSC group (47.2%) was similar to that of the FGSC group (49.9%). Subsequently, these zygotes developed to 2-cell embryos (Fig. 4e, Extended Data Fig. 11b). After the embryo transfer, 53 (male:female, 1.65:1.00) out of 342 (for iGSC group) or 51 (male:female, 1.13:1.00) out 355 (for FGSC group) were delivered as viable offspring with colored eyes (Fig. 4f, Extended Data Fig. 11c), indicating that the offspring were derived from C57BL/6 iGSC- or FGSC-derived oocytes, but not ICR oocytes among gonadal somatic cells. The offspring were confirmed for the presence of GFP transgenes by Southern blot analysis, and live imaging by a Lumazone imaging system (Fig. 4g, h). All of the obtained offspring grew up normally and were fertile with no difference between the two groups. After analysis of the methylation status, 10 offspring per group demonstrated no observably abnormal methylation patterns (Extended Data Table 1). Our findings provide a new strategy to investigate stem cell biology, biotechnology, and medicine.

## METHODS

### Mice

C57BL/6, pou5f1-GFP transgenic mice [CBA-Tg (pou5f1-EGFP) 2Mnn] (The Jackson Laboratory) or pou5f1/GFP transgenic mice^21^×C57BL/6 F1 hybrid mice were used in this study. Premature ovarian failure (POF) *Pten* (phosphatase and tensin homolog deleted on chromosome 10)^loxp/loxp^;*Gdf9-Cre* (*Gdf9* promoter-mediated Cre recombinase^+^) mice were produced and genotyped as described by Reddy *et al*^22,23^. POF (*Pten*^loxp/loxp^;*Gdf9*-Cre^+^) mice were used as recipients. *Pten*^*loxp/loxp*^mice (B6.129S4-*Pten*^tm1Hwu^) and *Gdf9-Cre* [Tg (*Gdf9-iCre*)] mice were purchased from The Jackson Laboratory. Animal experimentation was approved by the Institutional Animal Care and Use Committee of Shanghai and performed in accordance with the National Research Council Guide for Care and Use of Laboratory Animals.

### Isolation and culture of spermatogonial stem cells

Testes from 6-day-old pou5f1-GFP transgenic mice or pou5f1/GFP transgenic mice×C57BL/6 F1 hybrid mice were collected and decapsulated. Spermatogonial stem cells (SSCs) were isolated using methods described by Wu *et al*^24,25^ and Yuan *et al*^26^. The SSCs were purified by both magnetic activated cell sorting (MACS) with an anti-Thy-1 antibody and fluorescence activated cell sorting (FACS), according to the manufacturers’ instructions. SSCs were cultured on mitotically inactivated SIM mouse embryo derived thioguanine- and ouabain-resistant (STO) feeder cells (5 × 10^4^ cells/cm^2^; ATCC) in culture medium. For mitotic inactivation, STO cells were treated with 10 μg/ml mitomycin C (Sigma) for 2–3 hours. Mitomycin C-treated STO cells were washed with phosphate buffered saline (PBS) and transferred to 0.2% (w/v) gelatin-coated tissue culture plates. The SSC culture medium consisted of high glucose Dulbecco’s modified Eagle’s medium (DMEM; Life Technologies) supplemented with 10% fetal bovine serum (FBS; GIBCO), 2 mM L-glutamine (Sigma), 0.1 mM β-mercaptoethanol (Sigma), 1 mM nonessential amino acids (Life Technologies), 10 ng/ml glial cell line-derived neurotrophic factor (GDNF; R&D Systems), 10 ng/ml leukemia inhibitory factor (LIF; Chemicon), and 15 mg/l penicillin (Sigma). SSCs were cultured on STO feeders in 24-well plates with 500 μl culture medium per well. The medium was replaced every 1–2 days, and cells were subcultured at a split ratio of 1:1–3 by trypsinization every 3 days. All cultures were maintained at 37°C with 5% CO_2_.

### Isolation and purification of female germline stem cells

Ovaries were collected from 5-day-old pou5f1/GFP transgenic mice ×C57BL/6 F1 hybrid mice. Female germline stem cells (FGSCs) were isolated and purified using a method described elsewhere^27^. Briefly, dissected ovarian tissues were incubated in 1 mg/ml collagenase (type IV; Sigma) at 37°C with gentle agitation for 15–20 min. After washing, ovarian tissues were incubated in 0.05% trypsin and 1 mM EDTA at 37°C for 5–7 min. Sheep anti-mouse IgG magnetic beads (Dynal Biotech) were incubated with an anti-fragilis antibody (Abcam) for 30 min at room temperature. The magnetic bead/antibody mixture was incubated with the isolated cell suspension for another 30 min at room temperature. Then, the mixture of cells and magnetic beads was placed on a magnetic bead separator for 2–3 min, and the supernatant was removed. The fraction on the inner side of the eppendorf tube was collected and rinsed twice with PBS, resuspended in PBS, and further purified by FACS, in accordance with the manufacturers’ instructions. The purified FGSCs were placed in FGSC culture medium^27^ and cultured on mitotically inactivated STO feeder cells in 24-well plates at 37°C with 5% CO_2_.

### Preparation of ovarian tissue for analysis

Ovaries from recipient and control mice were fixed with 4% (w/v) paraformaldehyde (4°C, overnight) and dehydrated via a graded ethanol series. The tissues were vitrified in xylene, embedded in paraffin, sectioned (6 μm thickness), and then mounted on slides. Prior to immunofluorescence staining, the sections were dewaxed in xylene and rehydrated via a graded ethanol series. Sections were counterstained with hematoxylin.

### Immunofluorescence

After equilibration in PBS, tissue sections were digested with 0.125% trypsin for 10 min at 37°C and then washed in PBS twice. The sections were blocked in 10% goat serum at room temperature for 10 min and then incubated overnight at 4°C with appropriate primary antibodies. The primary antibodies used were mouse monoclonal anti-GFP (1:200 dilution; Abcam) and rabbit polyclonal anti-MVH (1:200; Abcam). After washing in PBS, the sections were incubated at 37°C for 30 min with TRITC-conjugated goat anti-rabbit IgG (1:200; Sino-American Biotechnology Co.) or fluorescein isothiocyanate (FITC)-conjugated goat anti-mouse IgG (1:200; Sino-American Biotechnology Co.) as appropriate. Sections were stained with 4′,6-diamidino-2-phenylindole (DAPI, 1:1000) at 37°C for 20 min, covered with mounting medium (glycerol:PBS, 3:1), and viewed under a Nikon Eclipse E600 microscope equipped with a Nikon Dxm 1200 digital camera using fluorescein optics for TRITC and FITC, and ultraviolet optics for DAPI or under a confocal microscope (FluoView™ FV1000).

Cultured germline stem cells were fixed with 4% paraformaldehyde in PBS at room temperature for 20 min. After fixation, the cells were permeabilized with 0.5% Triton X-100 for 30 min at room temperature for PLZF staining. The cells were incubated in blocking solution (10% normal goat or bovine serum in PBS, 10 min, 37°C), followed by rinsing and overnight incubation at 4°C with appropriate primary antibodies: rabbit polyclonal anti-MVH (1:200; Abcam), mouse monoclonal anti-GFP (1:200; Abcam), and anti-PLZF (1:150, Santa Cruz Biotechnology). After washing in PBS, the cells were incubated with TRITC-conjugated goat anti-rabbit IgG (1:200) or fluorescein isothiocyanate (FITC)-conjugated goat anti-mouse IgG (1:150) at 37°C for 30 min, rinsed, and then incubated with DAPI (1:1000) at 37°C for 20 min. Petri dishes were then covered with mounting medium (glycerol:PBS, 3:1) and viewed as described above.

### Karyotypic analysis

Karyotypic analysis was performed using standard protocols for mouse chromosome analysis. After culture for 3 days, SSCs were treated with culture medium containing colchicine (100 ng/ml; Sigma) for 3 hours, hypotonically treated with 75 mM KCl for 15 min at 37°C, immersed twice in methanol:acetic acid (3:1) for 30 min at −30°C, dried in airfor 3–4 days, digested with 0.025% trypsin, and then stained with Giemsa. To verify the chromosomal type of recipient mouse oocytes, karyotypic analysis of mature oocytes from recipients was performed as describedpreviously^28^. To collect mature oocytes, recipient mice were superovulated with 10 IU pregnant mare serum gonadotropin (PMSG; ProSpec-Tany) for 48 hours, followed by 10 IU human chorionic gonadotropin (hCG; ProSpec-Tany). These oocytes were hypotonically treated with 75 mM KCl at 37°C for 15 min and then fixed with two solutions consisting of methanol/acetic acid/water (5:1:2) for 5–10 min and methanol/acetic acid (3:1) for 15 min at room temperature. Fixed cells were mounted on slides and immediately exposed to steam from boiling water (90–100°C) for 30 sec to cause expansion of the cells, followed by drying at 37°C and Giemsa staining (Amresco)^28^.

### Embryonic stem cell culture

Embryonic stem cells (ESCs) were cultured with mouse embryonic fibroblasts in the presence of leukemia inhibitory factor (LIF; 1000 U/ml) in Glasgow modification of Eagle’s medium (GMEM; Invitrogen) containing 10% fetal calf serum. The medium was replaced every 1–2 days, and cells subcultured at a split ratio of 1:1–3 by trypsinization every 3 days. All cultures were maintained at 37°C with 5% CO_2_.

### Microarrays

Total RNA was extracted from cultured SSCs and ESCs using Trizol reagent (Invitrogen), in accordance with the manufacturer’s instructions. RNA was labeled using an Illumina labeling kit. An Illumina sentrix mouse WG-6 Beadchip (45281 transcripts) was used in this study. Microarray experiments, including RNA labeling, hybridization, washing, scanning, image analysis, normalization, and data processing, were performed by Shanghai Biotechnology Corporation using the Illumina manual. Three biological repeats were included in microarray experiments. Differentially expressed genes were identified by the Illumina system. The data were analyzed using GeneSpring GX 11 software. Hierarchical clustering of samples was performed by cluster 3.0 and TreeView software^29^.

### Transplantation

For injection into the ovary, SSCs were collected and transplanted into the ovaries of POF mice. For the positive control, FGSCs from pou5f1/GFP transgenic mice were also transplanted into ovaries of POF mice. Recipient mice were anesthetized by injection of pentobarbital sodium (45 mg/kg). Approximately 6 μl of a singlecell suspension containing 1 × 10^4^ cells or 6 μl PBS for the control was microinjected into the ovaries of recipientsas described elsewhere. In detail, after anesthetization of recipient mice for 20–30 min and disinfection of the abdominal surface using 75% ethanol, the recipient abdominal cavity was carefully opened. To expose and find the ovaries, the intestines were carefully moved away from the inside of the abdominal cavity. The Y-shaped uterus was located, and then following the uterus and oviduct until posterior to the kidneys, the ovaries were located caudal to the kidneys in the lower abdominal cavity. By gently holding an ovary with forceps without causing damage, the ovary was injected at 1–2 sites using a glass pipette with a 45 μm tip and mouth pipetting to carefully transplant the 6 µl single cell suspension of ∼1 × 10^4^ SSCs or FGSCs into each ovary. At 35 days after transplantation, recipients were mated with 8-week-old male mice.

### Reverse transcription-polymerase chain reaction and Southern blotting

Reverse transcription-polymerase chain reaction (RT-PCR), PCR, and Southern blotting were performed as described elsewhere. Twenty-five cycles of PCR were performed using Taq polymerase (Takara) with primer sets specific for each gene. The glyceraldehyde-3-phosphate dehydrogenase gene (*Gapdh*) was amplified in each sample as a loading control. PCR products were isolated, subcloned, and sequenced to confirm the gene sequence.

### Bisulfite genomic sequencing

Genomic DNA was extracted from SSCs, transplanted SSCs, ESCs, induced germline stem cells (iGSCs), and FGSCs. For bisulfite sequencing analysis of methylation, 500 ng genomic DNA was processed using an EZ DNA Methylation-Gold Kit™ (ZYMO Research), in accordance with the manufacturer’s instructions. The methylation status of imprinted genes was analyzed using specific primers (outside, 5′-GTTTTTTTGGTTATTGAAT-TTTAAAATTAGT-3′ and 5′-AAAAACCATTCCGTAAATACACAAATACCTA-3′, inside, 5′-TTAGTGTGGTTTATTATAGGAAGGTATAGAAGT-3′ and 5′-TAAACCTAAAATACTCAAAACTTTATCACAA-3′ for *H19*; 5′-GTG TAG AAT ATG GGG TTG TTT TAT ATT G-3′ and 5′-ATA ATA CAA CAA CAA CAA TAA CAA TC-3′ for *Rasgrf1*; 5′-GTA AAG TGA TTG GTT TTG TAT TTT TAA GTG-3′ and 5′-TTA ATT ACT CTC CTA CAA CTT TCC AAA TT-3′ for *Peg10*; 5′-TTA GTG GGG TAT TTT TAT TTG TAT GG-3′ and 5′-AAA TAT CCT AAA AAT ACA AAC TAC ACA A-3′ for *Igf2r;*outside, 5′-TATGTAATATGATATAGTTTAGAAATTAG-3′ and 5′-AATAAACCCAAATCTAAAATATTTTAATC-3′, inside, 5′-AATTTGTGTGATGTTTGTAATTATTTGG-3′ and 5′-ATAAAATACACTTTCACTACTAAAATCC-3′ for *Snrpn*). PCR products were sequenced and CpG islands were analyzed.

### PCR amplification of lineage-specific microsatellite loci

Genomic DNA was extracted from mouse tail tips or donor SSCs. DNA samples from donor SSCs, female recipients, mated males, and their corresponding offspring were analyzed by simple sequence length polymorphism (SSLP). Sequences for the primer pairs were designed according to the Mouse Genome Informatics website (http://www.informatics.jax.org/). Amplification of lineage-specific microsatellite DNA was performed in accordance with a previously described procedure^30^. PCR products were separated and analyzed by 3% agarose gel electrophoresis (Bio-Rad) and visualized by ethidium bromide staining.

### Flow cytometry and cell sorting

After MACS, the cells were suspended in PBS and subjected to flow cytometry to analyze and sort GFP-positive cells using a FACSAria II cell sorter equipped with BD software (Becton Dickinson).

### Quantitative reverse transcription-PCR analysis

Total RNA from cells was isolated using Trizol reagent. Complementary DNA was synthesized from 2 μg total RNA using a High Capacity cDNA Reverse Transcription Kit (Invitrogen). Primers were designed using Primer Premier Software (Primer Premier 5.0). Primer details are listed in Supplementary Table 4. *Gapdh* was amplified in each sample as an internal control. The mRNA level of each gene was normalized to *Gapdh* expression. The specificity of all quantitative real-time PCRs (qPCRs) was verified by a single peak in the melting curve. qPCRs were performed with a 7500 real-time PCR amplification system using SYBR Green PCR master mix (Applied Biosystems, UK). The relative levels of transcripts were calculated using the ΔΔCT method within the ABI 7500 System Software (V2.0.4). All gene expression levels were normalized to the internal standard gene, *Gapdh*. The means and standard error were calculated from triplicate measurements. Significance was determined using the Student’s *t*-test. A *P*-value of less than 0.05 was considered as significant, and a *P*-value of less than 0.01 was extremely significant.

### Single cell whole genome amplification and exome sequencing

Single cell whole genome amplification was performed on lysed single cells using a recently developed method named multiple annealing and looping based amplification cycles (MALBAC)^31^. In brief, amplification was initiated by primers, each with a 27 fixed and eight degenerate base hybridizing uniformly throughout the genome. Fragments with variable length at random starting positions were generated by polymerase extension for multiple cycles. All fragments were flanked by the 27 base-fixed sequence and their complementary sequences, and further amplified by PCR to about 1 µg for barcoded massively parallel sequencing on an Illumina HiSeq 2500 sequencing platform.

### Sry DNA in situ hybridization

We used a commercially available SRY DNA FISH kit (Mice SRY DNA biotin labelled POD and fluorescent FISH in situ hybridization double staining system, TBD Science), according to the manufacturer’s instructions. Briefly, sections were dewaxed with a graded series of ethanol, quenched in 3% H_2_O_2_ for 10 min at room temperature, and then washed twice with PBS. The sections were covered with SRY reagent B for 10 min at 37°C. After washing with PBS, the sections were incubated in Tris buffered saline (TBS) for 20 min at 95–100°C (pH 8.9) and then rinsed three times with cold TBS (5 min per rinse) and once with 0.2× saline sodium citrate (5 min per rinse) at 0 °C. The sections were incubated for 8 hours at 37°C with SRY reagent A and then washed three times with 2×saline sodium citrate at 37°C (3 min per rinse), three times with 0.2×saline sodium citrate (3 min per rinse) at 37 °C, and three times with TBS (2 min per rinse) at 37°C. The sections were covered with SRY reagent C for 45 min at 37°C. After washing in PBS, the sections were incubated at 37°C for 120 min with a fluorescein isothiocyanate (FITC)-conjugated mouse anti-digoxin monoclonal antibody, then incubated at 37°C for 120 min with DAPI. Finally, the sections were mounted in anti-fade mounting medium. Images were obtained using a Leica DMI3000 B microscope and Leica DFC550 digital camera.

### In situ high throughput chromosome conformation capture library generation using a low amount of cells

In situ high throughput chromosome conformation capture (Hi-C) assays were carried out according to the protocol with minor modifications^32-34^. Cells were fixed in a 1% final concentration of formaldehyde prior to 10 min incubation at room temperature. The reaction was quenched for 5 min by adding a 2.5 M glycine solution. Cells were pelleted twice (3000*g*, 4°C for 5 min), resuspended in ice-cold Hi-C lysis buffer for at least 15 min, and then washed once with 100 μl of 1× NEBuffer 2. The supernatant was discarded, and 1μl of 5% sodium dodecyl sulfate (SDS) was added to the remaining 9 μl solution. The pellet was gently mixed and incubated at 62°C for 10 min. After incubation, 9.5 μl water and 2.5 μl of 10% Triton X-100 were added to quench the SDS, and then the solution was incubated at 37°C for 30 min. Chromatin digestion was performed with Dpn II restriction enzyme (NEB, R0543M) at 37°C overnight and then inactivated for 20 min at 65 °C. To fill the overhangs generated by the Dpn II restriction enzyme, a master mix of 3.75 μl biotin-14-dATP (Life Technologies), 0.45 μl of 10 mM dCTP/dGTP/dTTP mix, and 1µl of 5 U/μl large DNA Polymerase I (NEB, M0210L) were added, followed by incubation at 24°Cfor 4 hours. The above biotin-labelled products were ligated by adding a master mix of 66.3 μl water, 12 μl of 10× NEB T4 DNA ligase buffer, 10 μl of 10% Triton X-100, 5μl of 10 mg/ml bovineserum albumin, and 2μl of 400 U/ml T4 DNA ligase, followed by incubation at 16°C for 20 hours and then inactivation at 75°C for 20 min. The samples were pelleted (3000*g*, 4°Cfor 5 min) and washed once with 100 μl of 10 mM Tris buffer. To remove biotin from unligated DNA ends, a master mix of 40 μl water, 5 μl of 10× NEBuffer 2.1, 0.125 μl of 10 mM dATP/dGTP, and 5 μl of 3,000 U/ml T4 DNA polymerase (NEB, M0203L) were added to the tube containing the DNA sample, followed by incubation at 20°C for 4 hours. The samples were pelleted (3000*g*, 4°Cfor 5 min) and resuspended in 50 μl of 10 mM Tris buffer. To digest the proteins, 2 μl of 20 mg/ml proteinase K (NEB, P8107S) was added, followed by incubation at 62°C for 18 hours and inactivation at 75°C for 30 min. The DNA was sheared to an average size of 400 bp (Covaris, M220) to perform the End Repair/dA-Tailing and Adaptor Ligation (NEB, E7337A) with a KAPA Hyper Prep Kit (KAPA, kk8502) and then processed by 3 μl of USER™ Enzyme (NEB, M5505L) at 37°C for 15 min to open up the loop. Biotin-labeled ligation products were isolated using MyOneStreptavidin T1 Dynabeads (Life Technologies, 65601) and then resuspended in 20 μl of 10 mM Tris buffer at 98°C for 10 min, and the supernatant was transferred to a fresh PCR tube. Hi-C DNA was amplified using Index Primers set 1 (NEB, E7335S). The Hi-C libraries were purified with AMPure XP beads (Beckman Coulter, A63881) and sequenced using an Illumina sequencing platform.

### Hi-C data processing, mapping, and ICE normalization

For Hi-C pair-end raw data, we first trimmed the adaptor sequences and low quality reads with BBmap (version 38.16). Then, we used HiCPro (version 2.7)^35^ to map, process, and perform iterative correction for normalization. Briefly, reads were independently aligned to the mouse reference genome (mm9) by the bowtie2 algorithm^36^. We discarded the uncut DNA reads, re-ligation reads, continuous reads, and PCR artifacts. We then used the unique mapped reads (MAPQ>10) to build the contact matrix. Valid read pairs were then binned at a specific resolution by dividing the genome into sequential bins of equal size. We generated the raw contact matrices at binning resolutions of 10, 20, 40, 100, and 200 kb. ICE^37^ normalization was applied to remove bias in the raw matrix, such as GC content, mappability, and effective fragment length in the Hi-C data.

### Validation of Hi-C data

The data reproducibility was confirmed by calculating Pearson’s correlation coefficient (PCC) between the two libraries. Briefly, the interaction frequency was generated for each pair of 40kb bins. For each possible interaction I_ij_ between two replicates, they were correlated by comparing each point interaction in the normalized interaction matrix. Considering that the interaction matrix was highly skewed toward proximal interactions, we restricted the correlation to a maximum distance of 2 Mb between points i and j. We used R to calculate Pearson’s correlation between two duplicates.

### Contact probability *p(s)*calculation

*P(s)* was calculated with normalized interaction matrices at a 40kb resolution, as describedpreviously^30^. *P(s)* calculations only considered intra interactions. Briefly, we divided the genome into 40kb bins. For each distance separated by 40, 80, 120, and 160 kb, we counted the number of interactions at corresponding distances. Then, we divided the number of interactions in each bin by the total number of possible region reads as *P(s)*. Furthermore, we normalized the sum of *P(s)* over the range of distances as 1. We used LOWESS fitting to construct the curve (log–log axis).

### Identification of A and B compartments

We used the R package (HiTC)^38^ pca.hic function to generate PC1 eigenvectors using 400kb normalized matrices with the following options: normPerExpected=TRUE, npc=1, for which a positive value indicated the A compartment, while a negative value indicated the B compartment. To investigate compartment switching, we defined switched bins only if PC1 eigenvectors changed in the same direction for two replicates.

### Identification of concordant genes with an A/B compartment switch

We used a previously described method with minor modifications to define genes with concordant changes in expression and compartment status^39^. Briefly, we calculated the covariance between the vector of the gene expression values (FPKM) and the vector of PC1 values for each gene across five cell types. The calculated covariance as a metric to quantitatively define “concordance” was used. We compared these observed covariance values with a random background distribution to calculate a P-value for the covariance for each gene. Then, we produced the background distribution by randomly shuffling the vector of FPKM for each gene and calculating the covariance between the PC1 values and random gene expression vector. A rank-based P-value could be calculated for observed covariance values with 1000 repeats for each gene. Concordant genes were defined as those with a P-value of <0.01.

### Direct induction of germline stem cells from SSCs

Knockdowns of specific genes were accomplished by small interfering RNAs (siRNAs) targeting *Plzf* and *Eed*. The interfering fragment was inserted downstream of the U6 promoter in a lentiviral vector (pLKD-CMV-G&PR-U6-shRNA) by molecular biological methods. At least four independent siRNAs were screened for knockdown efficiency against each target and the best siRNA target was selected (target Seq: CCAGGCATCTGATGACAAT for *Plzf*; GCAACAGAGTAACCTTATA for *Eed*). For *Stella, Zfp57, H19*, and *Rasgrf1* overexpression, cDNAs of candidate genes were inserted into the EcoRI and BamHI restriction sites of the overexpression plasmid (pHBLV-CMVIE-ZsGreen-T2A-puro). Lentivirus particles were generated by cotransfection of knockdown or overexpression plasmids and lentivirus packaging plasmids into HEK293T cells using transgene reagent. Enhancing buffer was added to the medium after 12 hours of transfection. Virus particles were harvested at 48 hours after transfection, and a standardized virus titer was obtained using HEK293T cells.

For lentivirus infection, 1×10^4^ SSCs, which were passaged for 2-3 times, were seeded in the well of a 48-well plate pre-coated with laminin and incubated with a 1:1 mixture of culture medium and lentivirus-concentrated solution (lentivirus titer: 1×10^9^TU/ml) containing 5μg/ml polybrene. After overnight infection, cells were re-plated onto puromycin-resistant STO feeder layers and cultured in SSC medium. At 12 hours after re-plating, the SSCs were incubated with a 1:1 mixture of culture medium and lentivirus-concentrated solution again. After overnight infection, the mixture was changed to fresh culture medium, and the cells were cultured for 12 hours. SSCs were then infected for a third time. After overnight infection, the mixture was changed to fresh culture medium, and the cells were cultured at 37°C with 5% CO_2_. At day 6, the cells were subcultured at a 1:1–2 split ratio, and 100 ng/ml puromycin was added to the FGSC culture medium to screen for puromycin-resistant iGSCs. After 72 hours, the surviving iGSCs were passaged and analyzed by qRT-PCR and western blotting.

### RNA-seq library generation and data analysis

Total RNA was extracted from 1–2×10^6^ cells using Trizol Reagent. The RNA quality was assessed using an Agilent Bioanalyzer 2100. RNA-Seq libraries were prepared using the KAPA Stranded mRNA-Seq kit, following the manufacturer’s instructions. After preparation, libraries were quantified using a Qubitfluorometer and sequenced with the HiSeq Platform (2×100 bp). All RNA-Seq data were trimmed and aligned to the mm9 reference genome using Hisat2 (version 4.8.2)^40^ with the default parameters. Gene expression as FPKM was calculated by Cufflinks (version 2.2.1)^41^ using the RefSeq database from the UCSC genome browser. Sequencing depth was normalized.

### GO term enrichment analysis

GO term enrichment analysis was performed using the DAVID tool (version 6.8)^42^, focusing on enriched biological processes (BP). The GO results were displayed by Cytoscape (version 3.5.1)^43^. For the Benjamin-corrected P-value, a threshold of <0.05 was used for significance.

### MeDIP-seq and bioinformatics

The DNA methylome assay was performed as described previously^44^. Briefly, genomic DNA (gDNA) was extracted and fragmented with Bioruptor (Connecticut, USA) into fragment sizes of 200–500 bp. Sonicated gDNA was used for end-repair and adaptor ligation. The adaptor-ligated gDNA was denatured and incubated with an antibody (Epigentek, A-1014) conjugated on Protein A+G Magnetic beads (Millipore, 16-663). Immunoprecipitated DNA was amplified by PCR and subjected to Illumina sequencing.

MeDIP and input raw sequencing reads were mapped using Bowtie2 (version 2.2.6) to the UCSC mm10 genome reference^36^. Duplicate reads were removed by samtools (version: 1.6-1). The normalized coverage was calculated by binning the unique tags in 1 kb bins, and the number of reads in each bin was normalized using reads per kilobase per million reads (RPKM). We identified the enriched MeDIP regions over the background with MACS (version 2.1.1) and default parameters^45^. Genome-wide pairwise correlation analysis of read depth in 1 kb bins was performed to evaluate DNA methylation patterns of SSCs, iGSCs, and FGSCs.

### Ovarian organoid generation and culture

Ovarian organoids were formed using a modified method described elsewhere^46^. Briefly, iGSCs, FGSCs (positive control), and SSCs (negative control) were purified by a FACS Aria II (BD Bioscience) and co-cultured with E12.5 female gonadal somatic cells in a 96-well U-bottom, low-binding culture plate (Thermo Fisher Scientific) for 2 days in GMEM supplemented with 15% Knockout serum replacement (Invitrogen), 1 μM retinoic acid, 2 mM L-glutamine (Sigma), 1 mM non-essential amino acids (Life Technologies), 2 mM L-glutamine (Sigma), 30 mg/ml pyruvate (Amresco), 50 mMβ-mercaptoethanol (Biotech), 30 mg/l penicillin (Amresco), and 75 mg/l streptomycin (Amresco). One thousand iGSCs, FGSCs or SSCs were 3D co-cultured with 3 × 10^4^ gonadal somatic cells. The co-cultures from 96-well U-bottom, low-binding culture plates were transferred onto transwell-COL membranes (Coaster) soaked in α-MEM--based medium, α-MEM supplemented with 2% FBS, 2 mM L-glutamine, 150 μM ascorbic acid (Sigma), 50 mM β-mercaptoethanol, 30 mg/lpenicillin, and 75 mg/l streptomycin. At 4 days of culture, the culture medium was changed to StemPro-34-based medium, StemPro-34 SFM (Life Technologies) supplemented with 10% FBS, 2 mM L-glutamine, 150 μM ascorbic acid, 50 mMβ-mercaptoethanol, 30 mg/l penicillin, and 75 mg/l streptomycin. From 7 to 10 days of culture, 600 nM ICI182780 was added to the StemPro-34-based medium. At 11 days of culture, the culture medium was changed to StemPro-34-based medium without ICI182780. After 21 days of culture, individual follicles were manually dissociated using sharpened tungsten needles.

### Follicle 3D culture

The single follicles were cultured on transwell-COL membranes with medium, α-MEM supplemented with 5% FBS, 2% polyvinylpyrrolidone (Sigma), 2 mM L-glutamine, 150 μM ascorbic acid, 50 mMβ-mercaptoethanol, 30 mg/l penicillin, 75 mg/l streptomycin, 30 mg/ml pyruvate (Amresco), 0.1 IU/ml follicle-stimulating hormone (FSH; MSD), 15 ng /ml BMP15, and 15 ng/ml GDF9 (R&D Systems). At 2 days of culture, the culture medium was changed to medium without BMP15 and GDF9, and then follicles were incubated in 0.1% Type IV Collagenase (Invitrogen) for 5 min. After washing with α-MEM supplemented with 5% FBS several times, the follicles were cultured in medium without BMP15 and GDF9. After 14 days of culture, cumulus-oocyte complexes grown on the membrane were picked up by a fine glass capillary.

### In vitro maturation, in vitro fertilization, and embryo transfer

The cumulus-oocyte complexes were cultured with α-MEM containing 5% FBS, 30 mg/ml pyruvate (Amresco), 0.1 IU /ml FSH, 4 ng/ml EGF, 1.2 IU/ml hCG (gonadotropin, ASKA), 4 ng/ml bFGF, 30 mg/l penicillin, 75 mg/l streptomycin. After 17-20 hours of culture, mature oocytes with expanded cumulus cells were fertilized in HTF medium (SAGE) by sperm. Embryos developed to the 2-cell stage were transferred into the oviducts of pseudopregnant females at 0.5 day post-coitum.

## Data Availability

Original data of Hi-C have been deposited in the Gene Expression Omnibus database (accession number: 135104). Original data of RNA-Seq have been deposited in the Gene Expression Omnibus database (accession number: 134727). Original data of MeDIP-Seq have been deposited in the Gene Expression Omnibus database (accession number: 134640). Original data of Microarrays have been deposited in the Gene Expression Omnibus database (accession number: GSE38776). All other relevant data are available from the corresponding author upon request.

## Acknowledgements

This work was supported by the National Key Research and Development Program of China (2018YFC1003501, 2017YFA0504201), National Nature Science Foundation of China (81720108017), the National Major Scientific Instruments and Equipment Development Project, National Nature Science Foundation of China (61827814).

## Author contributions

H.L., X.L. and G.G.T. conducted all the major experiments, data analysis and wrote the manuscript; D.L. performed embryo transfer; C. H. carried out in situ Hi-C library generation using a low amount of cells; X.D. and W.X. were responsible for karyotype analysis; L.H, Y.Y., and H.W. were responsible for immunofluorescence and histological analysis of ovarian tissue; Q.L., A.J.C. and J.X. conducted Gdf9-Cre^+^and GFP transgenic mice study; X.Z. carried out MeDIP-seq and bioinformatics; J.W. initiated and supervised the entire project, conducted SSC and FGSC transplantation, analyzed data and wrote the manuscript.

## Competing interests

The authors declare no competing interests.

### Materials & Correspondence

Correspondence and requests for materials should be addressed to J.W.

