## Supplemental file for "Offspring production of ovarian organoids derived from spermatogonial stem cells by chromatin reorganization"

**Extended Data Figure Legends and Tables**

**Extended Data Fig. 1: Characterization of spermatogonial stem cells** **from neonatal mice. a,** Representative example of spermatogonial stem cell (SSC) purification by fluorescence-activated cell sorting (FACS). **b,** Merge of bright field and fluorescence images of SSCs after purification by FACS. **c,** Merge of bright field and fluorescence microscopies of cultured SSCs. **d, e,** Cytogenetic analysis by G-band staining showing that SSCs possessed a normal karyotype (40, XY). **f,** Gene expression profiles of SSCs. M, 100 bp DNA marker; lane 1, SSCs; lane 2, Embryonic stem cells (ESCs); lane 3, mock-transcribed SSC RNA samples. **g-j,** Cultured SSCs were positive for EGFP (**g**) and MVH (mouse vasa homologue, expressed exclusively in germ cells) (**h**) staining. Cells were counterstained with 4',6-diamidino-2-phenylindole (DAPI) (**i**). **k, l,** Global gene expression profiles of SSCs and ESCs. (**k**) Scatter plots of gene expression values in SSCs versus ESCs. Average expression levels of each gene (a dot on scatter plots) were calculated from three independent experiments. Genes whose expression level was at least two times greater in SSCs are shown in red, and genes whose expression level was at least two times greater in ESCs are shown in green. (**l**) Clustering analysis of some pluripotency-related genes and germ cell markers between SSCs and ESCs. **m-p,** DMR methylation pattern of *H19* (**m**)*, Igf2r* (**n**)*, Rasgrf1* (**o**)*,* and *Peg10* (**p**) regions. DNA methylation levels were analyzed by bisulfite genomic sequencing. Black circles represent methylated cytosine-guanine sites (CpGs), and white circles represent unmethylated CpGs. The percentage of methylated CpG sites is shown beside the map. Scale bars: 50 μm (**b, c**), 30 μm (**g-j**).

**Extended Data Fig. 2: Offspring derived from Female germline stem cells**. **a,** Example of female germ cells isolated by magnetic activated cell sorting (MACS) with an anti-fragilis antibody. **b,** Representative examples of female germline stem cell (FGSC) purification by FACS. **c,** Representative morphology of ovaries from recipients of FGSC transplantation. **d,** GFP-positive (green, arrows) oocytes in recipient ovaries at 6–8 weeks after transplantation of Pou5f1/GFP transgenic FGSCs. **e,** Oocytes (arrows) in a wild-type ovary did not show a GFP signal. **f,** Example of offspring from premature ovarian failure (POF) recipient mice transplanted with Pou5f1/GFP transgenic FGSCs. **g,** Example of Southern blotting (see above). Lanes 1, 2, and 6, transgenic mice; lanes 3–5, wild-type mice. Scale bars, 10μm (**a**) and 100 μm (**c-e**).

**Extended Data Fig. 3: Methylation status and gene expression dynamics of transplanted SSCs in recipient ovaries. a-d**, Methylation status of *H19* (**a, c**) and *Peg10* (**b, d**) DMRs in SSCs at 0 and 2 hours, and 2, 3, 4, 5, 6, 9, 12, and 15 days after transplantation. DNA methylation levels were analyzed by bisulfite sequencing. Black circles represent methylated cytosine-guanine sites (CpGs), and white circles represent unmethylatedCpGs. The percentage of methylated CpG sites is shown in (**c)** and **(d)**. **e,** Gene expression dynamics in transplanted SSCs at 0 and 2 hours, 2, 3, 4, 5, 6, 9, and 15 days after transplantation.

**Extended Data Fig. 4: Validation high throughput chromosome conformation capture data of Pou5f1-GFP transgenic SSCs and FGSCs, and overview of whole genome interaction frequency heat maps in germline stem cells. a**, Correlation between high throughput chromosome conformation capture (Hi-C) replicates for SSCs and FGSCs according to the normalized interaction frequency at a 400kb resolution. R indicates Pearson’s correlation coefficient. **b,** Overview of whole genome interaction frequency heat maps in SSCs and FGSCs. **c,** Hieratical clustering of PC1 values for the A/B compartment status in SSCs and FGSCs.

**Extended Data Fig. 5: Effects of specific gene expression levels examined by qRT-PCR and western blot analyses**. Data are presented as the mean ± SD of three independent experiments. **P< 0.01 compared with the control group.

**Extended Data Fig. 6: Screening the critical imprinted genes and transcription factor genes required for SSC conversion. a,** Methylation analysis at paternally imprinted loci (*H19* and *Rasgrf1*) and maternally imprinted loci (*Igf2r*, *Snrpn*, and *Peg10*) in SSCs, iGSCs (induced by 6Gs, *Stella, H19, Zfp57, Rasgrf1, Plzf,* and *Zfp42*), and FGSCs. **b,** Representative merged bright field and fluorescence images of ovarian organoids. I, Ovarian organoids formed by FGSCs co-cultured with somatic cells of gonads. II, Ovarian organoids formed by iGSCs induced by 6Gs co-cultured with somatic cells of gonads. III, Withdrawal of *Rasgrf1* from 6Gs, and ovarian organoids formed by iGSCs induced by remaining in 5Gs (*Stella, H19, Zfp57, Plzf,* and *Zfp42*) co-cultured with somatic cells of gonads. IV, Withdrawal of *Zfp42* from 5Gs, and ovarian organoids formed by iGSCs induced by remaining in 4Gs (*Stella, H19, Zfp57*,and *Plzf*) co-cultured with somatic cells of gonads. V–VIII, Removal of *Stella* (V), *H19* (VI)*, Zfp57* (VII), or *Plzf* (VIII) from 4Gs failed to form ovarian organoids. **c,** Methylation analysis at paternally imprinted loci (*H19* and *Rasgrf1*) and maternally imprinted loci (*Igf2r*, *Snrpn* and *Peg10*) in iGSCs induced by 4Gs. Scale bars: 100 μm.

**Extended Data Fig. 7: Characterization of induced germline stem cells derived from SSCs in vitro**. **a,** Morphology and immunofluorescence detection of MVH and GFP in cultured induced germline stem cells (iGSCs) and FGSCs. I, II, Representative morphology of cultured iGSCs under bright field (I) and fluorescence (II) microscopies. III, IV, Representative view of cultured FGSCs under bright field (III) and fluorescence (IV) microscopies. V–VII, Cultured iGSCs were positive for EGFP (V) and MVH (VI) staining. Cells were counterstained with DAPI (VII). VIII, Merge of EGFP, MVH, and DAPI staining. IX–XI, Cultured FGSCs were positive for EGFP (IX) and MVH staining (X). Cells were counterstained with DAPI (XI). XII, Merge of EGFP, MVH, and DAPI staining. **b,** RT-PCR analysis of germ cell or germline stem cell markers in SSCs, iGSCs, and FGSCs. Lane M, 250bp DNA marker; lane 1, SSCs, lane 2, iGSCs, lane 3, FGSCs, lane 4, STO, lane 5, no template control. **c,** Genomic view of the DNA methylation pattern defined by MeDIP-seq in the IGV (Integrative Genomics Viewer) genome browser and DNA methylation status of imprinting gene *Igf2r* in SSCs, iGSCs, and FGSCs. **d,** Pairwise correlation comparison of genome-wide DNA methylation among SSCs, iGSCs, and FGSCs. R values (Pearson correlation coefficient) were used to compare the significant correlation both within and between groups and is represented by a color scale. **e,** Heat map showing expression profiles of genes among SSCs, iGSCs, and FGSCs. The maps were based on the expression values of all expressed genes detected by high-throughput sequencing. The color scale indicates the expression values. The intensity increases from green to red. Each column represents one sample, and each row represents a transcript. **f,** RNA-seq read density over imprinting gene *Igf2r* in SSCs, iGSCs, and FGSCs.Scale bars, 50 μm (**a** I-IV), 20 μm (**a** V-XII).

**Extended Data Fig. 8: RNA-Seq quality control. a,** Fast-QC data showing the position-specific sequencing quality in each replicated sample of SSCs, iGSCs, and FGSCs. **b,** Correlation plots of each replicated sample of SSCs, iGSCs, and FGSCs.

**Extended Data Fig. 9: Validation Hi-C data of iGSCs and overview of whole genome interaction frequency heat maps in iGSCs. a,** Correlation between Hi-C replicates of iGSCs according to the normalized interaction frequency at a 400kb resolution. R indicates Pearson’s correlation coefficient. **b,** Hieratical clustering of PC1 values for the A/B compartment status in SSCs, iGSCs, and FGSCs. **c,** Overview of whole genome interaction frequency heat maps in iGSCs.

**Extended Data Fig. 10: Gene expression dynamics during oogenesis *in vitro* determined by qRT-PCR.** Relative mRNA expression of genes in oogenesis of iGSCs (red) and FGSCs (blue) are shown. 1, GSCs, 2, immature oocytes, and 3, MII oocytes.

**Extended Data Fig. 11: The percentage of oocyte maturation and fertilization and number of offspring derived from iGSCs or FGSCs**. **a,** The percentage of mature cumulus-oocyte complexes derived from iGSCs and FGSCs. **b,** The percentage of fertilization of iGSC- and FGSC-derived MII oocytes. **c,** Number of offspring derived from iGSCs or FGSCs per litter.

**Table 1 The DNA methlytion analysis of offspring derived from iGSCs and FGSCs**

| Source of offspring mice | Number of  selected offspring | The DNA methylation levels (%) | |
| --- | --- | --- | --- |
|  |  | H19 | Peg10 |
| **Wide type** |  |  |  |
|  | WT-1 | 49.55 | 48.51 |
|  | WT-2 | 50.77 | 48.96 |
|  | WT-3 | 49.22 | 49.03 |
| **FGSCs** |  |  |  |
|  | F-1 | 58.87 | 50.75 |
|  | F-2 | 51.22 | 51.53 |
|  | F-3 | 50.34 | 49.41 |
|  | F-4 | 49.38 | 51.62 |
|  | F-5 | 48.34 | 48.93 |
|  | F-6 | 49.44 | 49.19 |
|  | F-7 | 51.79 | 48.50 |
|  | F-8 | 50.43 | 51.77 |
|  | F-9 | 38.54 | 49.67 |
|  | F-10 | 51.46 | 48.68 |
| **IGSCs** |  |  |  |
|  | I-1 | 49.17 | 51.13 |
|  | I-2 | 51.32 | 48.54 |
|  | I-3 | 51.08 | 49.16 |
|  | I-4 | 49.16 | 51.83 |
|  | I-5 | 49.00 | 48.77 |
|  | I-6 | 49.68 | 50.98 |
|  | I-7 | 50.33 | 50.75 |
|  | I-8 | 51.71 | 49.66 |
|  | I-9 | 48.52 | 49.53 |
|  | I-10 | 49.31 | 49.19 |
