## Supplementary figures and images for "Offspring production of ovarian organoids derived from spermatogonial stem cells by chromatin reorganization"

### Supplemental Fig. 1

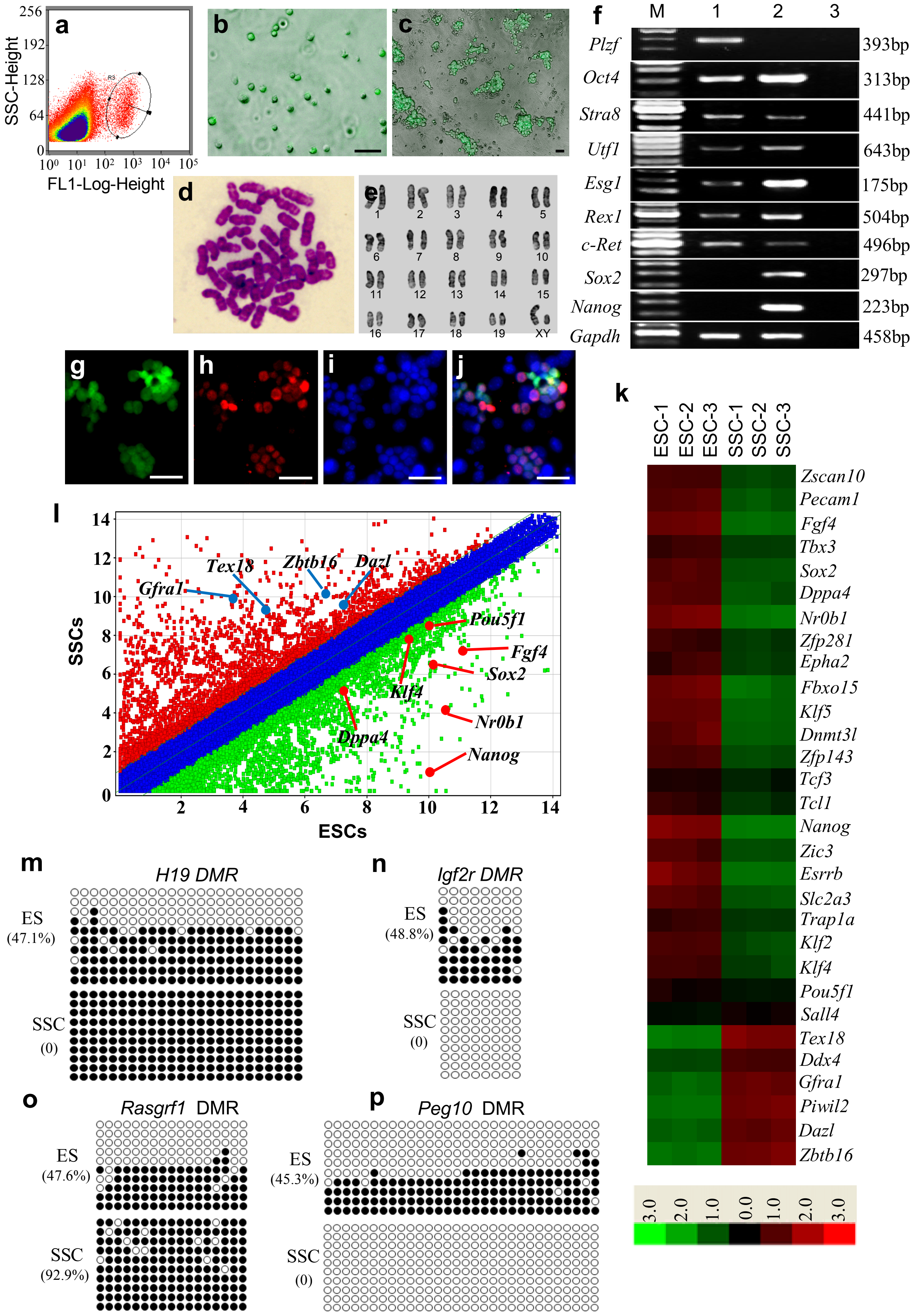

### Supplemental Fig. 2

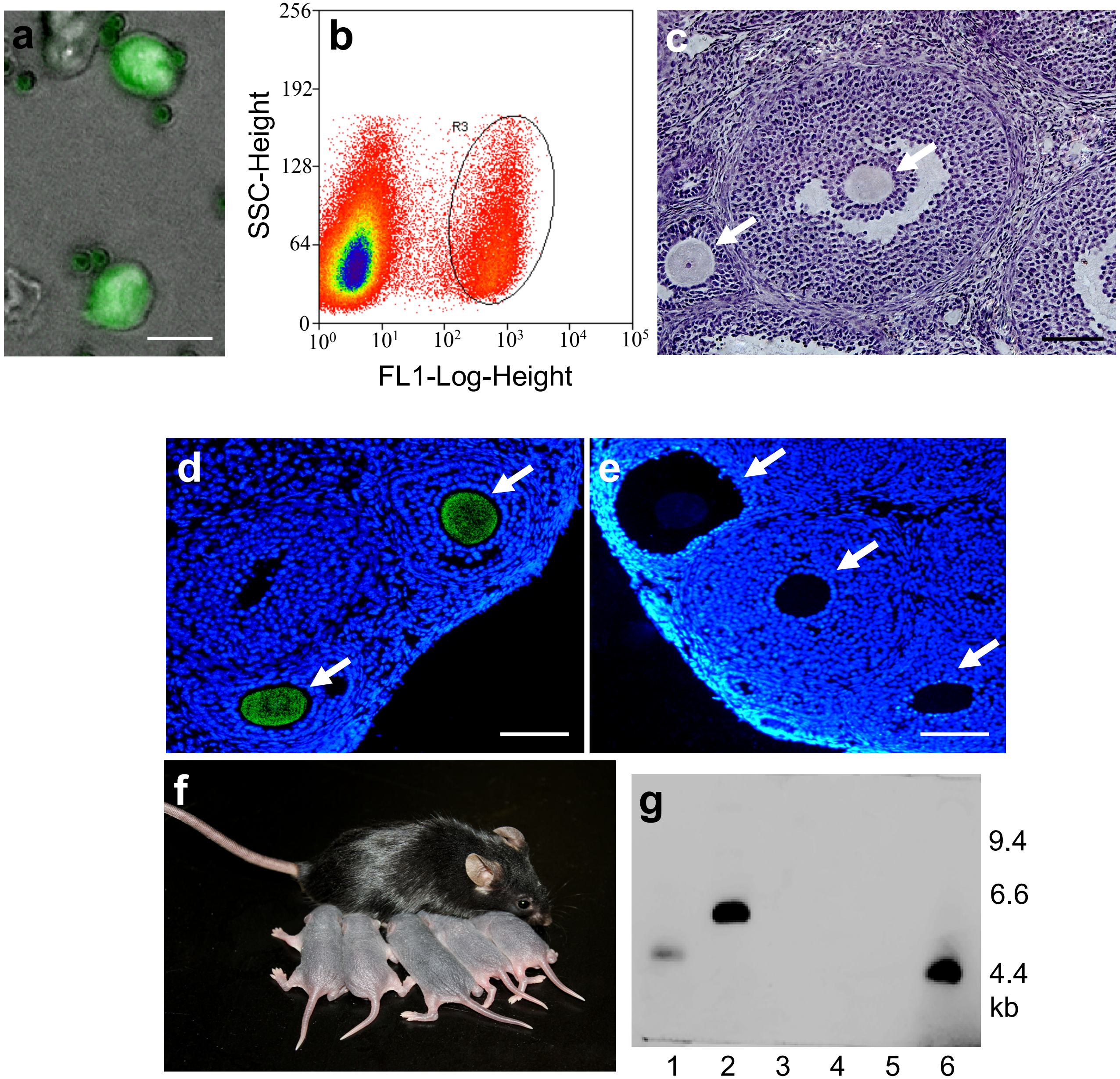

### Supplemental Fig. 3

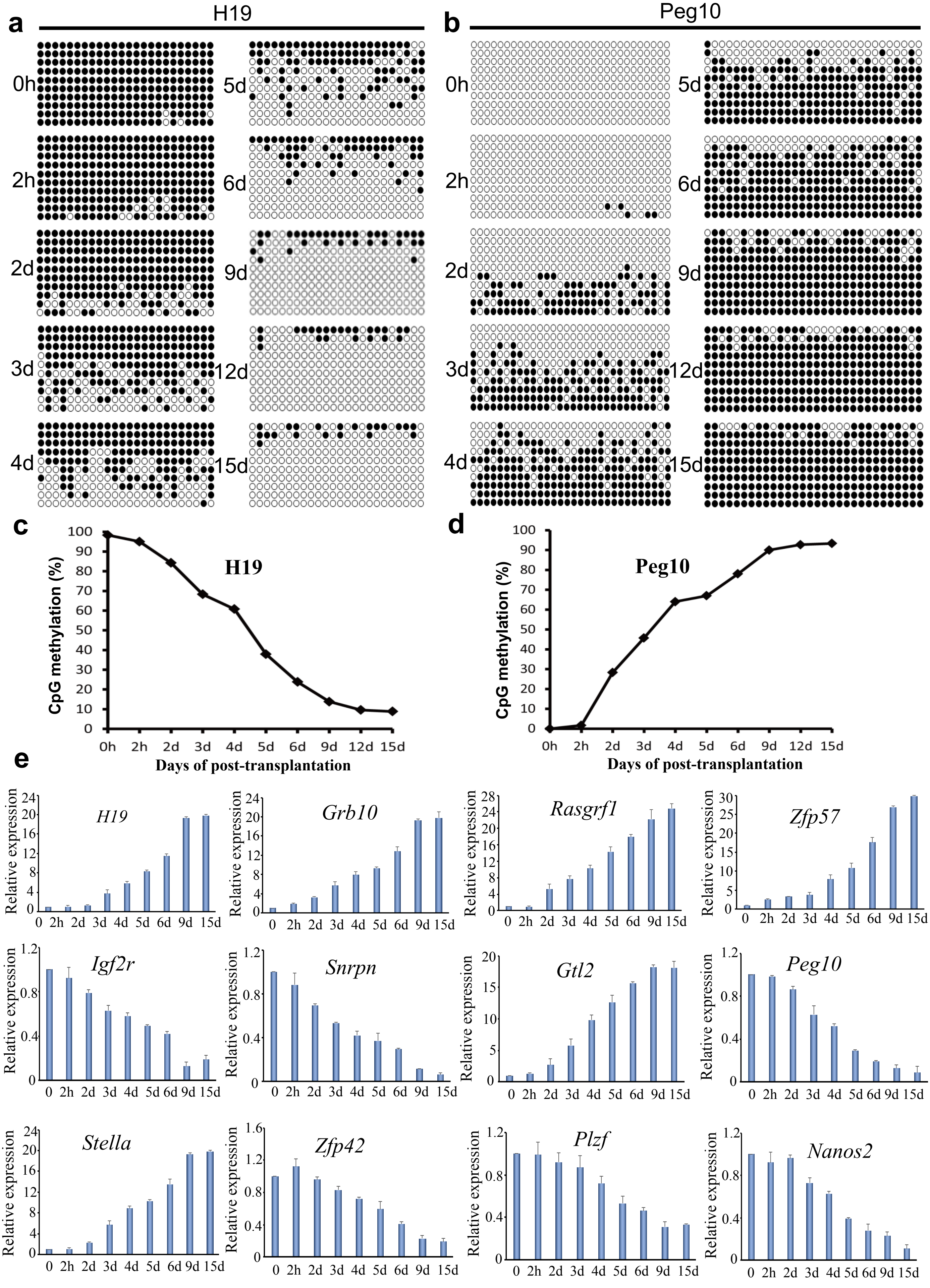

### Supplemental Fig. 4

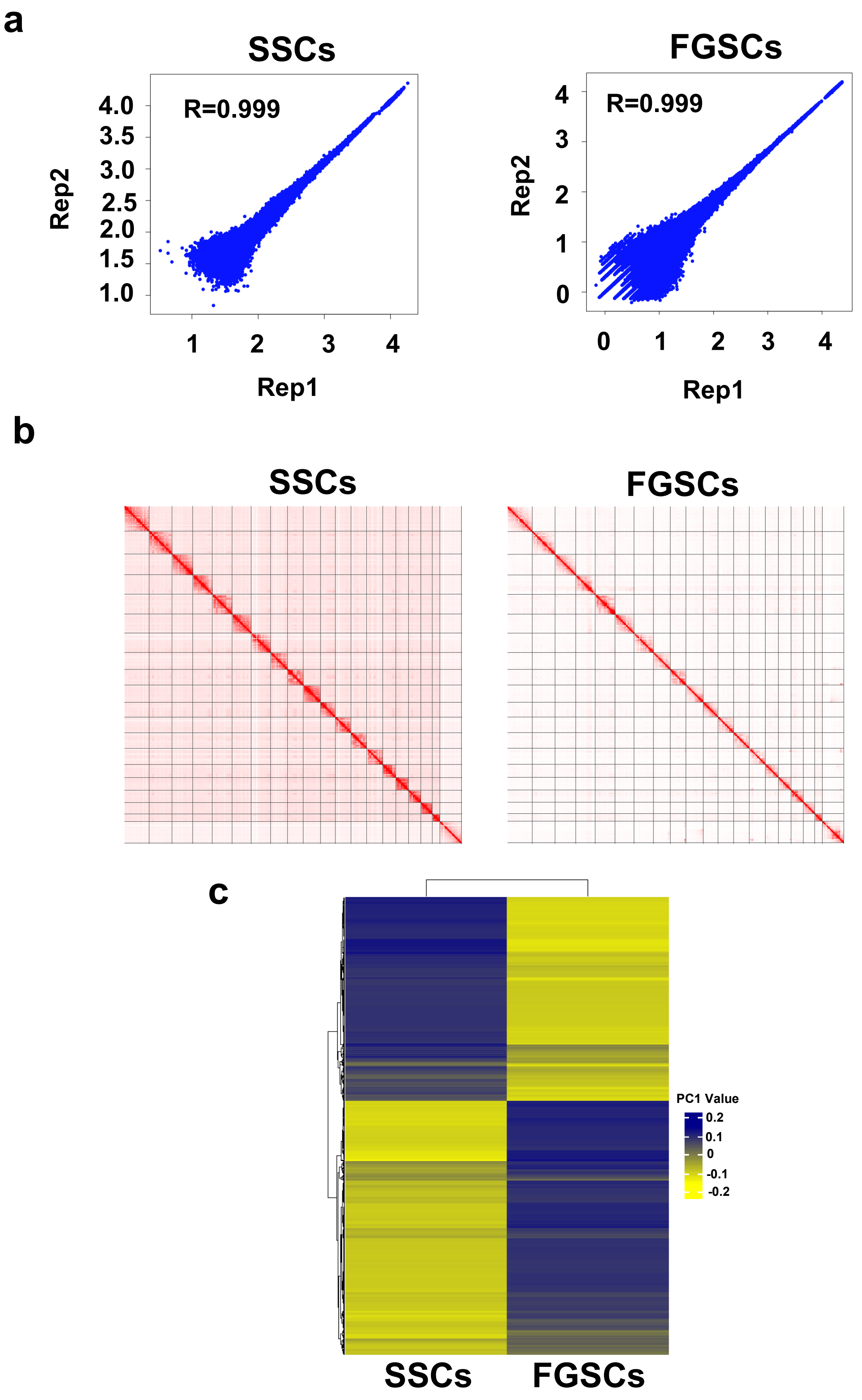

### Supplemental Fig. 5

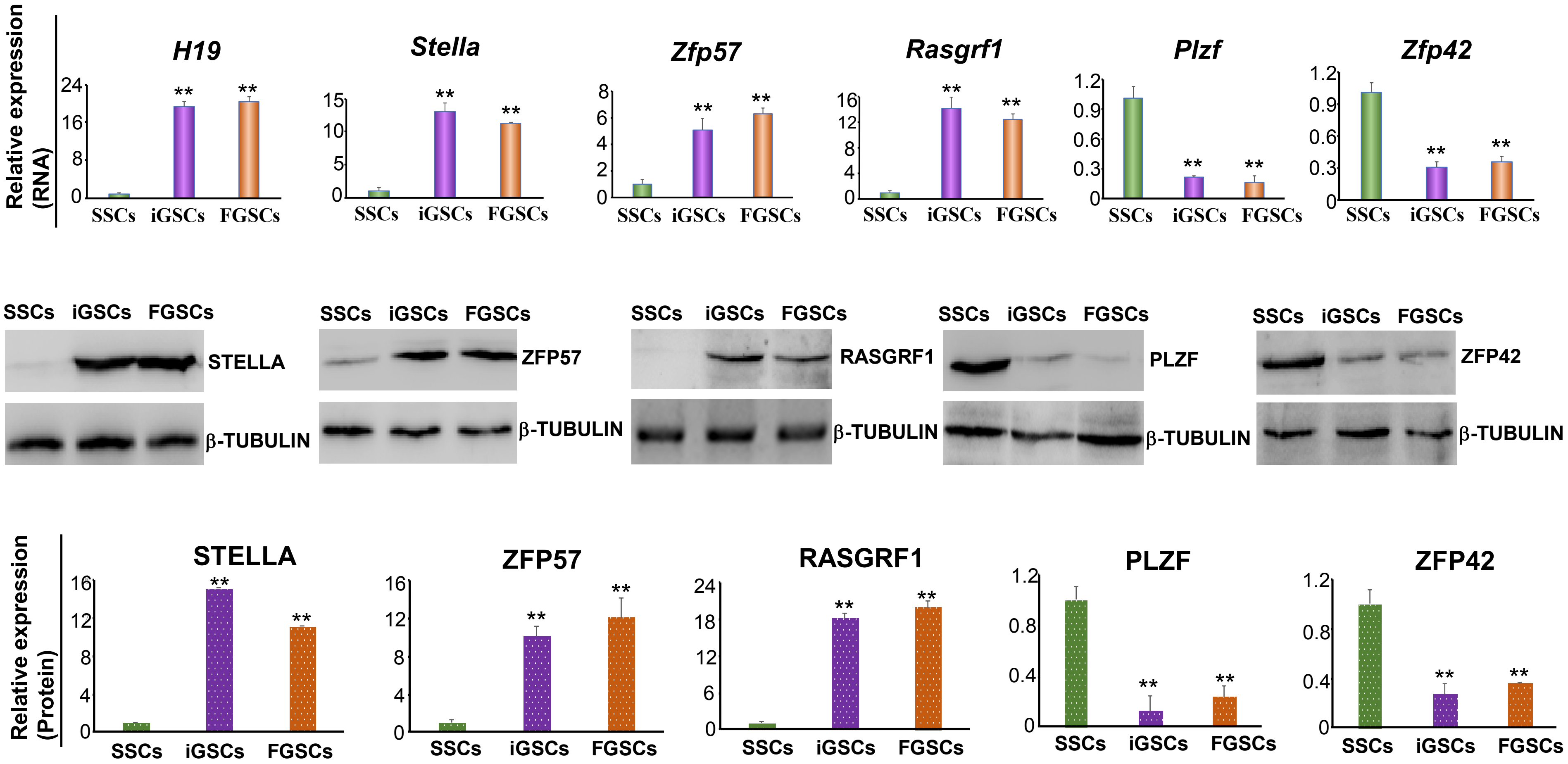

### Supplemental Fig. 6

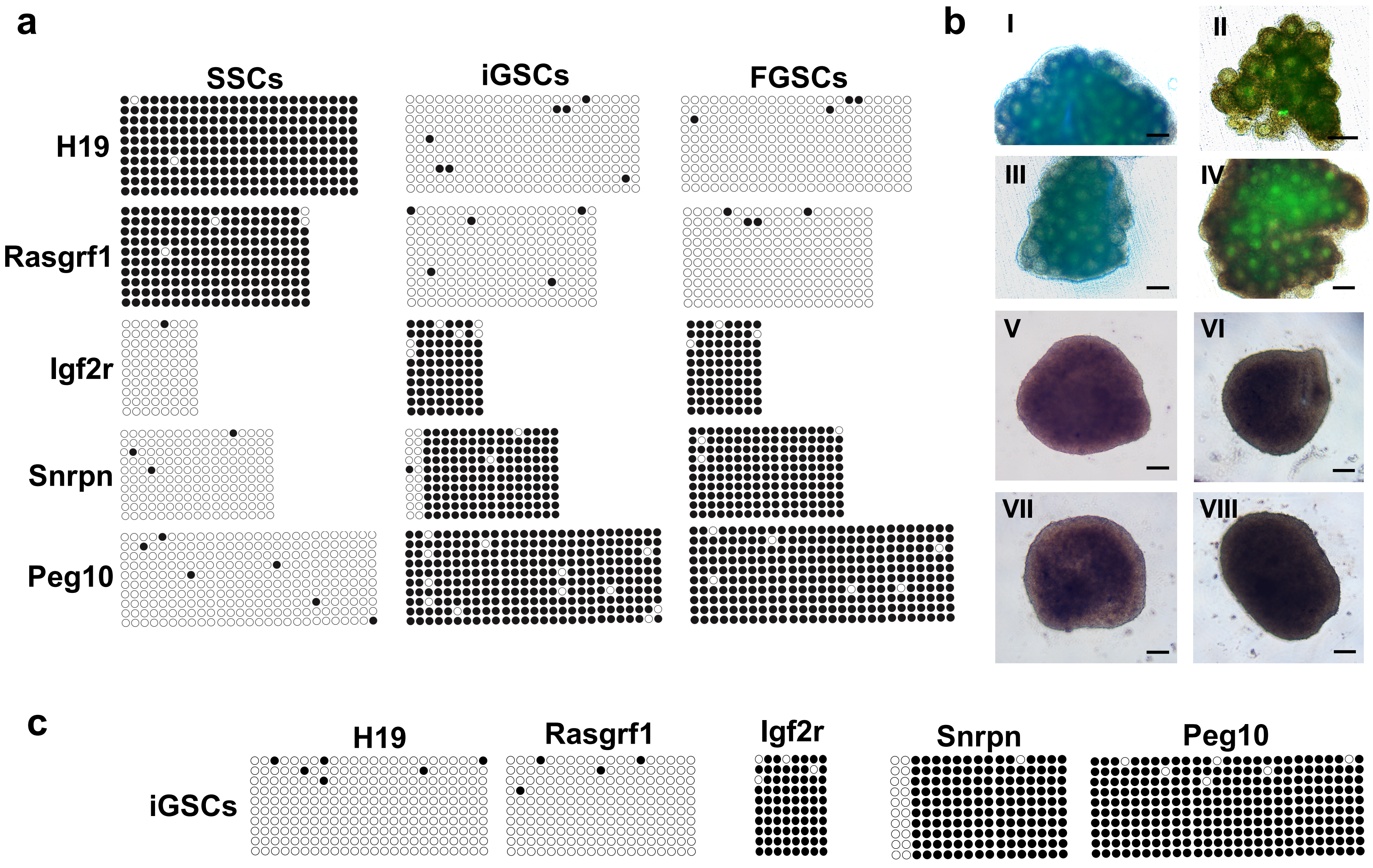

### Supplemental Fig. 7

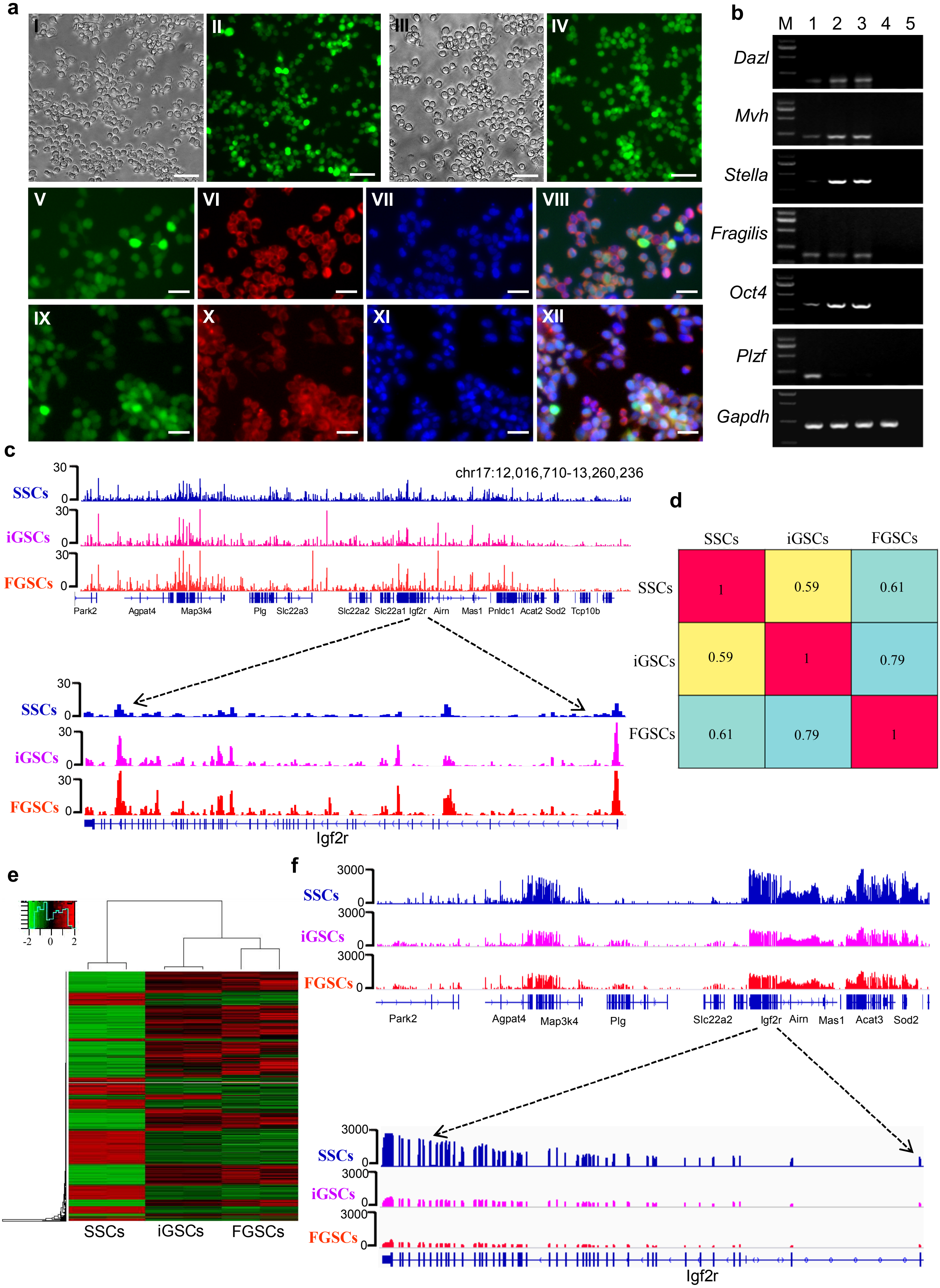

### Supplemental Fig. 8

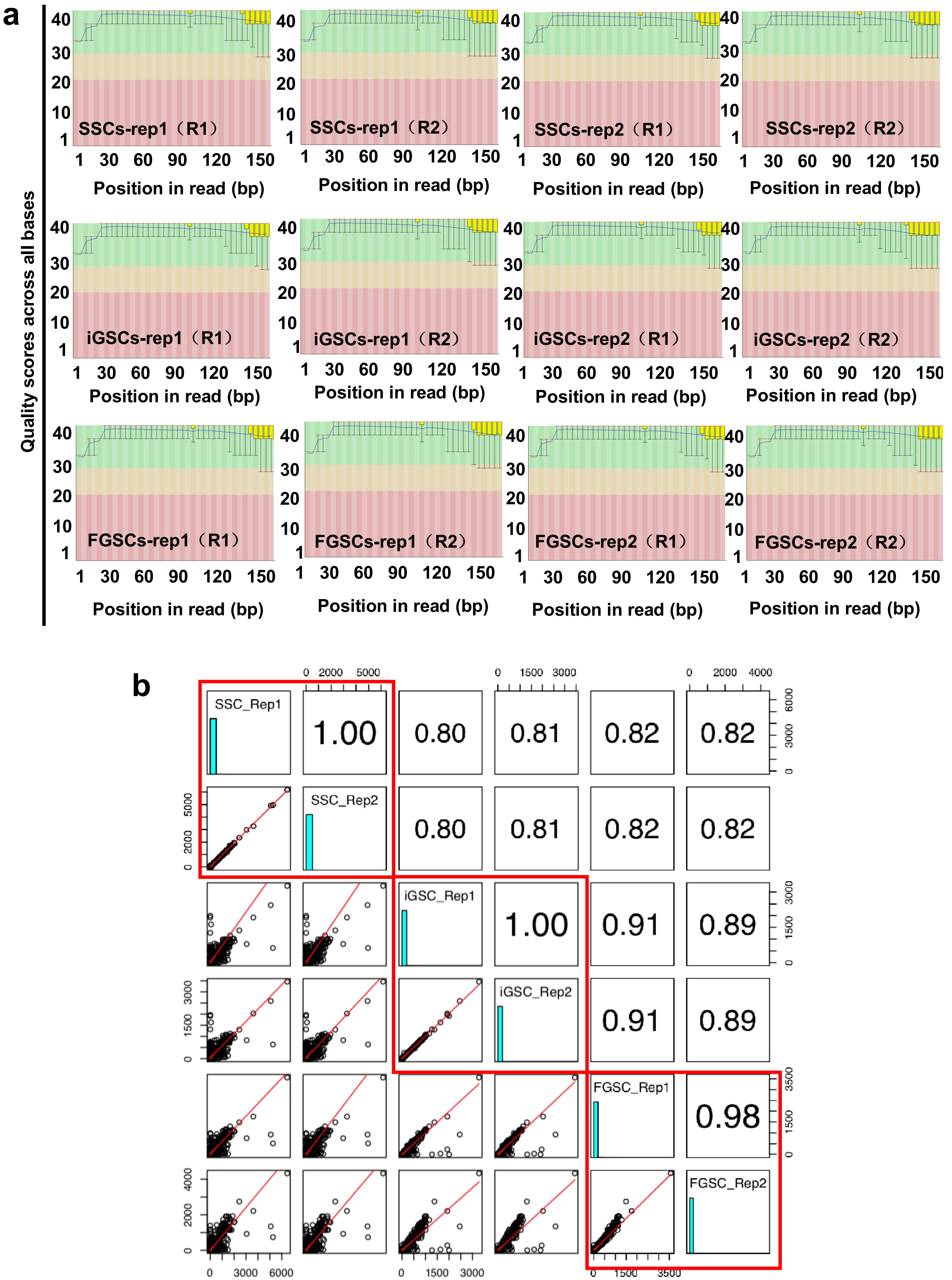

### Supplemental Fig. 9

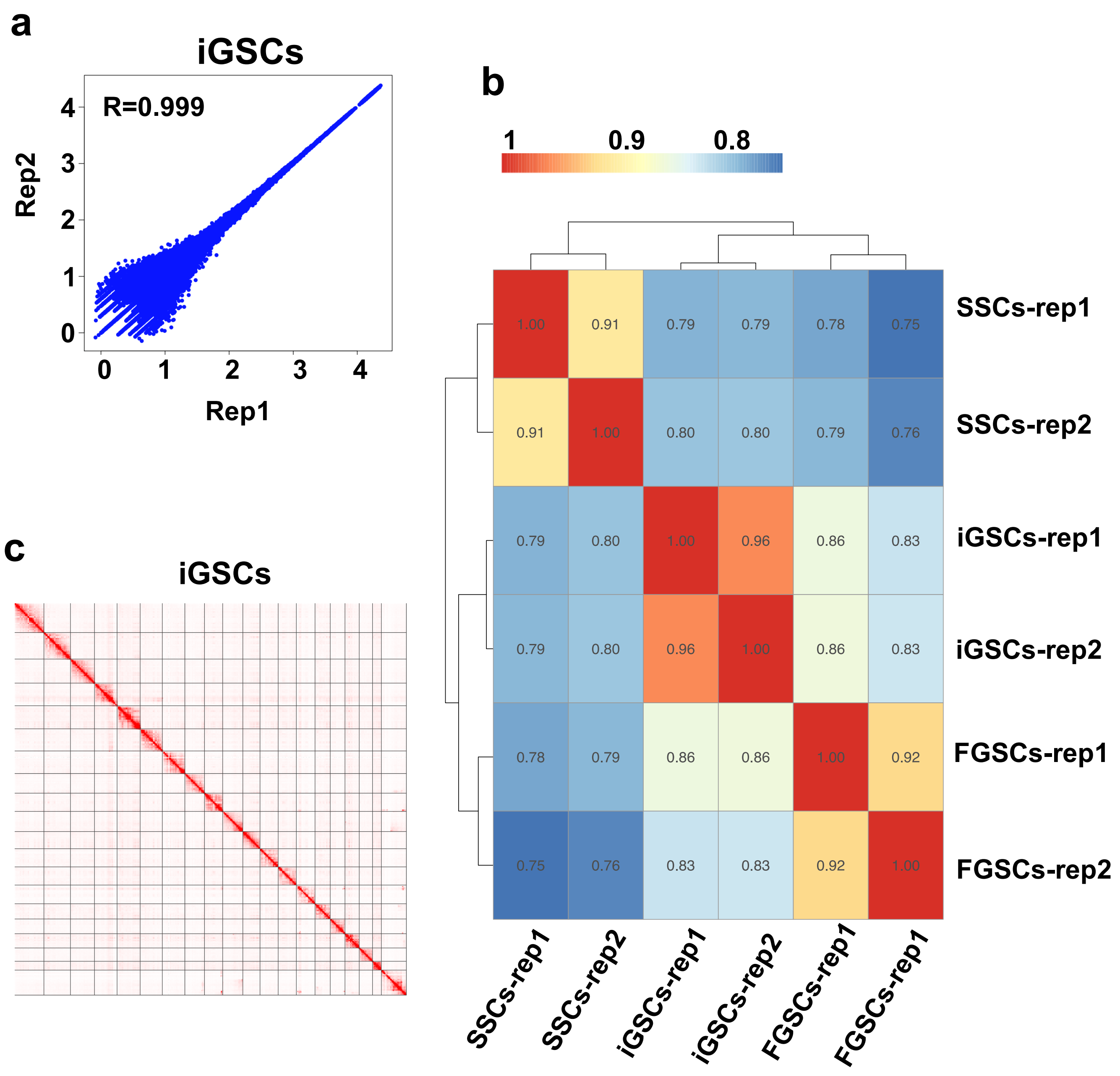

### Supplemental Fig. 10

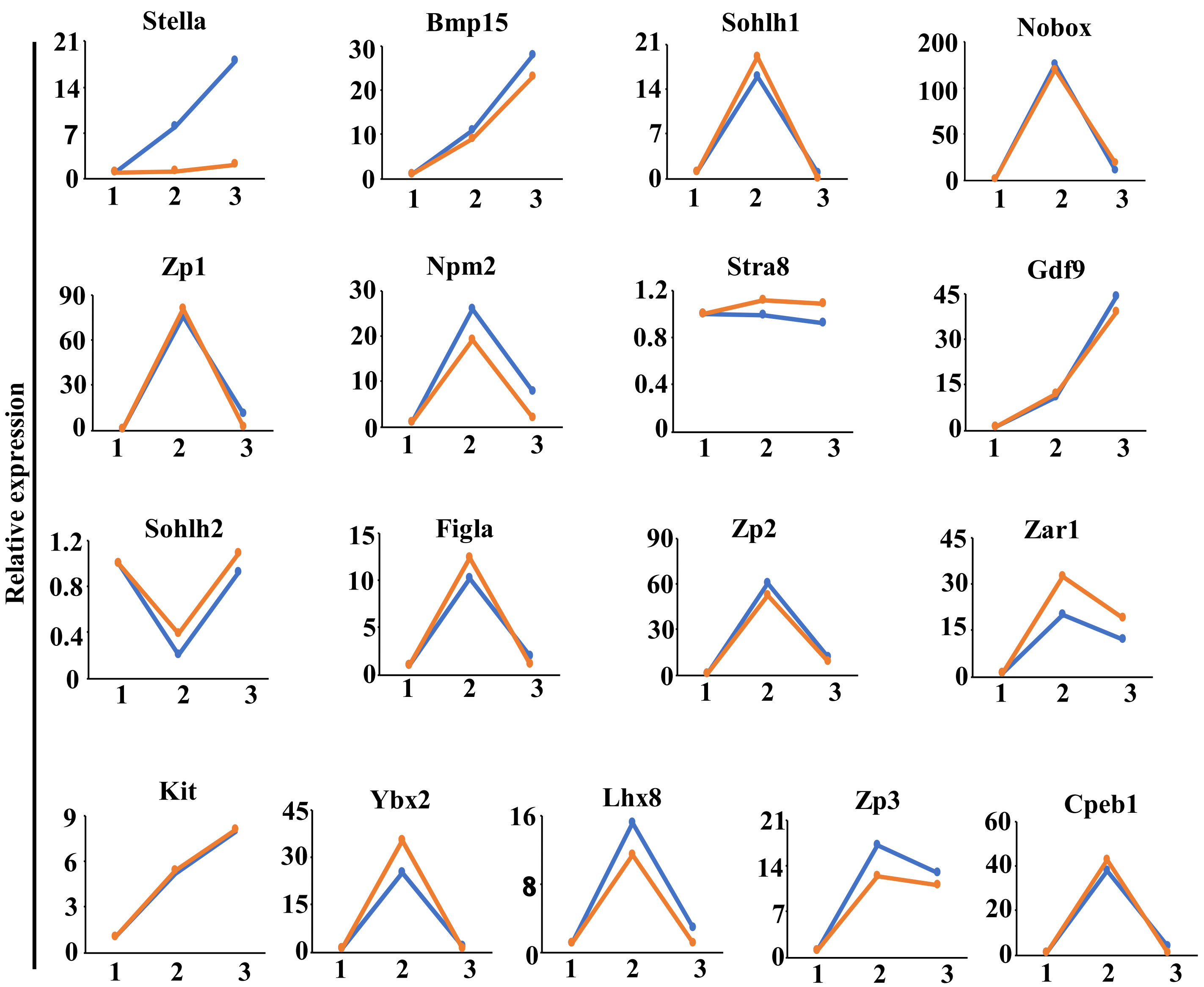

### Supplemental Fig. 11

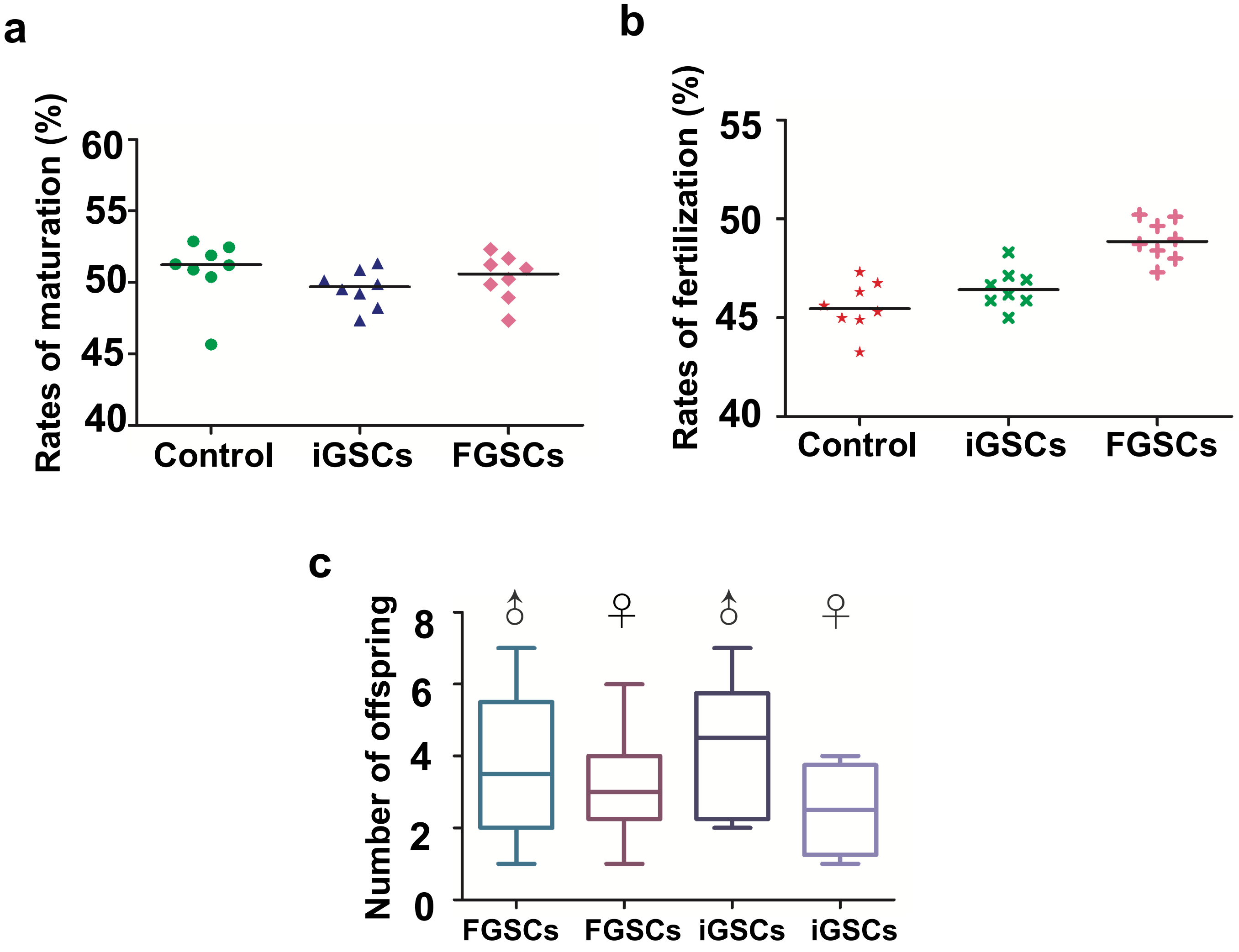
