## Supplemental Table 4 for "Offspring production of ovarian organoids derived from spermatogonial stem cells by chromatin reorganization"

**Supplement Table 4. Details regarding the RT-PCR or qRT-PCR analysis of cells and tissues**

| Gene | Product  Size(bp) | Primer Sequence (5’-3’) |
| --- | --- | --- |
| **Primers for gene expression dynamic during oogenesis *in vivo* (Fig. 2B)** | | |
| Plzf | 393 | F：CACCAACCTTTCTTCTCCGGG |
|  |  | R：CCGTGTAGGCGTACTCCAGG |
| Mvh | 193 | F: GCTTCATCAGATATTGGCGAGT |
|  |  | R: GCTTGGAAAACCCTCTGCTT |
| Stra8 | 173 | F: ACAACCTAAGGAAGGCAGTTTAC |
|  |  | R: GACCTCCTCTAAGCTGTTGGG |
| Sycp3 | 206 | F: AGCCAGTAACCAGAAAATTGAGC |
|  |  | R: CCACTGCTGCAACACATTCATA |
| Zp1 | 164 | F: CCCTGAGATTGGGTCAGCG |
|  |  | R: AGAGCAGTTATTCACCTCAAACC |
| Zp3 | 186 | F: ATGGCGTCAAGCTATTTCCTC |
|  |  | R: CGTGCCAAAAAGGTCTCTACT |
| Gapdh | 123 | F: AGGTCGGTGTGAACGGATTTG |
|  |  | R: TGTAGACCATGTAGTTGAGGTCA |
| **Primers for gene expression dynamic in transplanted SSCs (Fig. 3E)** | | |
| H19 | 106 | F: GAACAGAAGCATTCTAGGCTGG |
|  |  | R: TTCTAAGTGAATTACGGTGGGTG |
| Grb10 | 181 | F: CCTGCCAAGCATGATGTCAAA |
|  |  | R: CCAGGCACCTCTCTAATCCCA |
| Rasgrf1 | 71 | F: GCCAGAAGACTTGACAACGCT |
|  |  | R: TCAATCTACAGGGATGGTGGAAG |
| Zfp57 | 122 | F: ATGGCAGCTAGGAAACAGTCT |
|  |  | R: TGGTAAAGGGTCTTCTGTGTAGA |
| Igf2r | 194 | F: GGGAAGCTGTTGACTCCAAAA |
|  |  | R: GCAGCCCATAGTGGTGTTGAA |
| Snrpn | 156 | F: TGCTACGTGGGGAGAACTTG |
|  |  | R: CCTGGGGAATAGGTACACCTG |
| Gtl2 | 71 | F: TCCTCACCTCCAATTTCCCCT |
|  |  | R: GAGCGAGAGCCGTTCGATG |
| Peg10 | 162 | F: TGCTTGCACAGAGCTACAGTC |
|  |  | R: AGTTTGGGATAGGGGCTGCT |
| Stella | 130 | F: GACCCAATGAAGGACCCTGAA |
|  |  | R: GCTTGACACCGGGGTTTAG |
| Zfp42 | 161 | F: GGAGGAAATAGGTAGAGCGCA |
|  |  | R: AGTGAGGCGATCCTGCTTTC |
| Plzf | 150 | F: CTGCGGAAAACGGTTCCTG |
|  |  | R: GTGCCAGTATGGGTCTGTCT |
| Nanos2 | 183 | F: CTGCAAGCACAATGGGGAGT |
|  |  | R: CGTCGGTAGAGAGACTGCTG |
| Gapdh | 123 | F: AGGTCGGTGTGAACGGATTTG |
|  |  | R: TGTAGACCATGTAGTTGAGGTCA |
| **Primers for RT-PCR analysis of germ cell or germline stem cell markers (Fig. 4B)** | | |
| Dazl | 170 | F: ATGTCTGCCACAACTTCTGAG |
|  |  | R: CTGATTTCGGTTTCATCCATCCT |
| Mvh | 193 | F: GCTTCATCAGATATTGGCGAGT |
|  |  | R: GCTTGGAAAACCCTCTGCTT |
| Stella | 386 | F: ATCGCCATGGAGGAACCATC |
|  |  | R: AATGGCTCACTGTCCCGTTC |
| Fragilis | 183 | F: GCCTATGCCTACTCCGTGAA |
|  |  | R: AGTGTGAAGGTTTTGAGCGTT |
| Oct4 | 313 | F: GGCGTTCTCTTTGGAAAGGTGTTC |
|  |  | R: CTCGAACCACATCCTTCTCT |
| Plzf | 150 | F: CTGCGGAAAACGGTTCCTG |
|  |  | R: GTGCCAGTATGGGTCTGTCT |
| Gapdh | 222 | F: CAGGAGAGTGTTTCCTCGTCC |
|  |  | R: TTCCCATTCTCGGCCTTGAC |
| **Primers for gene expression dynamic during oogenesis *in vitro* (Fig. 6D)** | | |
| Stella | 130 | F: GACCCAATGAAGGACCCTGAA |
|  |  | R: GCTTGACACCGGGGTTTAG |
| Bmp15 | 100 | F: TCCTTGCTGACGACCCTACAT |
|  |  | R: TACCTCAGGGGATAGCCTTGG |
| Sohlh1 | 181 | F: CGGGCCAATGAGGATTACAGA |
|  |  | R: TCCTGCGTTCTCTCTCGCT |
| Nobox | 189 | F: ATGGAACCTACGGAGAAGCTC |
|  |  | R: CTCAGAGGTCTTCGACAGTGG |
| Zp1 | 164 | F: CCCTGAGATTGGGTCAGCG |
|  |  | R: AGAGCAGTTATTCACCTCAAACC |
| Npm2 | 162 | F: GTGACCGAAACCACAGCAAAA |
|  |  | R: CACACGGTTCACCTCCTCTT |
| Stra8 | 173 | F: ACAACCTAAGGAAGGCAGTTTAC |
|  |  | R: GACCTCCTCTAAGCTGTTGGG |
| Gdf9 | 116 | F: TCTTAGTAGCCTTAGCTCTCAGG |
|  |  | R: TGTCAGTCCCATCTACAGGCA |
| Sohlh2 | 154 | F: GGGCAGGGCAGAGTAAATCTT |
|  |  | R: CAAACGAGTTAGCAGCCAAAAG |
| Figla | 238 | F: CCGCCATCTGTAGGCTCAAG |
|  |  | R: ACACAGCCGAGTATCTGTATGTA |
| Zp2 | 111 | F: GTGGCAGAGGAAAGCATCTGT |
|  |  | R: GACTGAGGAAGGCTTACTGAGT |
| Zar1 | 136 | F: TCGGTGCAGTGTTCACTCG |
|  |  | R: CTACGGTCTGCCAGGATCG |
| Kit | 90 | F: GCCACGTCTCAGCCATCTG |
|  |  | R: GTCGCCAGCTTCAACTATTAACT |
| Ybx2 | 102 | F: GGAGTTTGATGTCGTGGAAGG |
|  |  | R: CGTCGATTAGGGGCATAGCG |
| Lhx8 | 159 | F: TCAGAGAGTGGTTACGGTCAC |
|  |  | R: CTGCTCGTCACATACCAGCTC |
| Zp3 | 186 | F: ATGGCGTCAAGCTATTTCCTC |
|  |  | R: CGTGCCAAAAAGGTCTCTACT |
| Cpeb1 | 229 | F: AAGGATTGCTGGGACAACCAA |
|  |  | R: GGCCACGGGGAGATTCTTG |
| Gapdh | 123 | F: AGGTCGGTGTGAACGGATTTG |
|  |  | R: TGTAGACCATGTAGTTGAGGTCA |
| **Primers for gene expression profiles of SSCs (Fig. S1F)** | | |
| Gapdh | 458 | F: GTCCCGTAGACAAAATGGTGA |
|  |  | R: TGCATTGCTGACAATCTTGAG |
| Oct4 | 313 | F: GGCGTTCTCTTTGGAAAGGTGTTC |
|  |  | R: CTCGAACCACATCCTTCTCT |
| Rex-1 | 504 | F: CACCATCCGGGATGAAAGTGAGAT |
|  |  | R: ACCAGAAAATGTCGCTTTAGTTTC |
| Esg-1 | 175 | F: GCCGTGCGTGGTGGATAAGC |
|  |  | R: GCCAAACAGATATTTCAGCACCAGC |
| Stra8 | 441 | F: TCACAGCCTCAAAGTGGCAGG |
|  |  | R: GCAACAGAGTGGAGGAGGAGT |
| Utf1 | 643 | F: GATGTCCCGGTGACTACGTCT |
|  |  | R: TCGGGGAGGATTCGAAGGTAT |
| c-Ret | 496 | F: TGGAAGCAGGAGCCAGACA |
|  |  | R: TGCTCTAATCCGCTTCTCCTG |
| Sox-2 | 297 | F: TAGAGCTAGACTCCGGGCGATGA |
|  |  | R: TTGCCTTAAACAAGACCACGAAA |
| Nanog | 223 | F: CAGGAGTTTGAGGGTAGCTC |
|  |  | R: CGGTTCATCATGGTACAGTC |
| **Primers for effects of specific gene expression levels examined by qRT-PCR (Fig. S4)** | | |
| H19 | 106 | F: GAACAGAAGCATTCTAGGCTGG |
|  |  | R: TTCTAAGTGAATTACGGTGGGTG |
| Stella | 130 | F: GACCCAATGAAGGACCCTGAA |
|  |  | R: GCTTGACACCGGGGTTTAG |
| Zfp57 | 122 | F: ATGGCAGCTAGGAAACAGTCT |
|  |  | R: TGGTAAAGGGTCTTCTGTGTAGA |
| Rasgrf1 | 71 | F: GCCAGAAGACTTGACAACGCT |
|  |  | R: TCAATCTACAGGGATGGTGGAAG |
| Plzf | 150 | F: CTGCGGAAAACGGTTCCTG |
|  |  | R: GTGCCAGTATGGGTCTGTCT |
| Zfp42 | 161 | F: GGAGGAAATAGGTAGAGCGCA |
|  |  | R: AGTGAGGCGATCCTGCTTTC |
| Gapdh | 123 | F: AGGTCGGTGTGAACGGATTTG |
|  |  | R: TGTAGACCATGTAGTTGAGGTCA |

F, forward primer; R, reverse primer
